# Impairment of neuronal activity occurs at the early stages of the aggregation cascade of Aβ_1-42_ and mutant Tau

**DOI:** 10.1101/2025.07.21.665894

**Authors:** Franziska Hirsch, Nino Läubli, Anushree Kelkar, Mira Sleiman, Yvonne Woitzat, Christian Gallrein, Vanessa Gumz, Gurleen Kaur Kalsi, Ana Fernandez-Villegas, Gabriele Kaminski Schierle, Janine Kirstein

## Abstract

Alzheimer’s disease (AD) is a progressive neurodegenerative disease that is characterized by the accumulation of amyloid-β (Aβ) plaques and neurofibrillary Tau tangles, ultimately leading to brain atrophy and death. To elucidate the relationship between the aberrant folding and aggregation of Aβ and mutant Tau and neuronal function, we monitored neuronal activity in *C. elegans* AD models across age. Our findings reveal that expression of both Aβ and Tau lead to significant reductions in neuronal activity and function in young adult animals preceding the accumulation of amyloid aggregates. Notably, Aβ expression and aggregation in muscle tissue produced comparable detrimental effects on neuronal activity as its expression in neurons, suggesting that proteotoxic stress in muscle can influence neuronal function. This may occur through the propagation of Aβ from muscle to neurons or through retrograde signaling pathways. Further, our new sub-stoichiometrically labeled Tau strains highlight that Tau^P301L,V337M^ has a significant impact on neuronal activity throughout aging. These results enhance our understanding of the early functional effects of amyloid aggregation in Alzheimer’s disease.

## Introduction

Alzheimer’s disease (AD) is the most common form of dementia and characterized by a progressive decrease in neuronal activity and severe neurodegeneration. The neuropathological hallmarks of AD are the accumulation of amyloid-β (Aβ) plaques and neurofibrillary Tau tangles. However, in the early presymptomatic stages of the disease, neuronal hyperactivity of cortical and hippocampal brain areas was observed in sporadic and familial forms of AD ^1,2^. Several factors could lead to the hyperexcitability of selected neurons and neuronal circuits in AD including perturbed Ca^2+^ homeostasis, NMDA receptor activation, Aβ and Tau themselves as well as genetic risk factors such as apolipoprotein E (APOE) and presenilin 1 and 2 (PSEN1 and PSEN2) ^3^. Thus far, it is not understood how hyperexcitability eventually leads to reduced neuronal activity. As neuronal activity promotes the production and secretion of Aβ by e.g. activating endocytosis ^4^, the hyperexcitability in the early stages of AD could be a pathogenic driver of the disease. However, in contrast to murine Aβ models, Tau^P301L^ expressing mice did not show a hyperexcitability and were marked by a loss of excitatory neuron function ^5,6^. Additionally, transgenic mouse models that express mutant Tau^P301S^ showed impaired synaptic function in the hippocampus and mutant Tau^P301S,G272V^ mice displayed axonopathy without overt neuronal loss before fibrillary Tau tangles were formed ^7,8^. Nevertheless, despite an increasing number of studies on AD pathology in diverse model systems, it is poorly understood how the expression and aberrant folding and amyloid fibrils formation of Aβ and mutant Tau lead to neurodegeneration.

In this study, we aimed to elucidate how the expression and aggregation of human Aβ and Tau in *C. elegans* are linked to neuronal activity as aging and pathology progress. For that, we used *C. elegans* Aβ and Tau models, analyzed neuronal activity, function, and organismal physiology over the course of Aβ and Tau aggregation. Neuronal activity was measured using a genetically encoded Ca^2+^ sensor, GCaMP6m. To facilitate a rapid and reliable assessment of neuronal activity of our AD *C. elegans* models, we established a microfluidic device that permits the trapping and immobilization of living nematodes in a non-invasive manner.

We have further established a new sub-stoichiometrically fluorescently tagged neuronal Tau pathology model in *C. elegans* that carries two patient derived point mutations, P301L and V337M ^9,10^. The pan-neuronal expression of Tau^P301L,V337M^ led to age-dependent aggregation, cell-cell propagation, and severe proteotoxicity on cellular and organismal level. Using our previously established Aβ _11_ and the novel Tau model, we found that both AD-associated amyloid proteins lead to a significant decrease of neuronal activity and function already in young adults before an accumulation of amyloid aggregates. Importantly, neuronal activity is equally impaired when the amyloid protein is expressed in a peripheral tissue, suggesting a retrograde signaling of the proteotoxic challenge to the neurons.

## Results

### Microfluidic devices permit the study of neuronal activity in living animals by quantifying GCaMP6m fluorescence intensity levels

In this study, we aimed to assess neuronal activity in Alzheimer’s disease models that express the Aβ_1-42_ peptide as well as wild-type Tau (Tau^WT^) and mutant Tau (Tau^P301L,V337M^) to correlate the aggregation propensity and proteotoxicity of Aβ_1-42_ and Tau with their effect on neuronal activity and function.

Neuronal activity can be analyzed using calcium-dependent fluorescence measurements by applying genetically encoded calcium indicators such as GCaMP ^12^. As neuronal action potentials lead to an influx of Ca^2+^ ions through voltage-gated Ca^2+^ channels, this results in a rapid increase in intracellular Ca^2+^ concentrations that are sensed by the GCaMP sensor and used as a proxy for neuronal activity ^13^. We have generated a *C. elegans* neuronal GCaMP strain that expresses GCaMP6m pan-neuronally (Fig. 1A). These imaging analyses are challenging in living animals that can move out of the imaging area or out of focus. Applying anesthetics on the other hand will affect their neuronal activity. To restrict the movement of the animals while recording the neuronal GCaMP fluorescence of the whole animal, we established a microfluidic device that traps the nematodes (Fig. 1B and Fig. S1). This allows a non-invasive quantification of GCaMP fluorescence in living animals.

**Figure 1.**
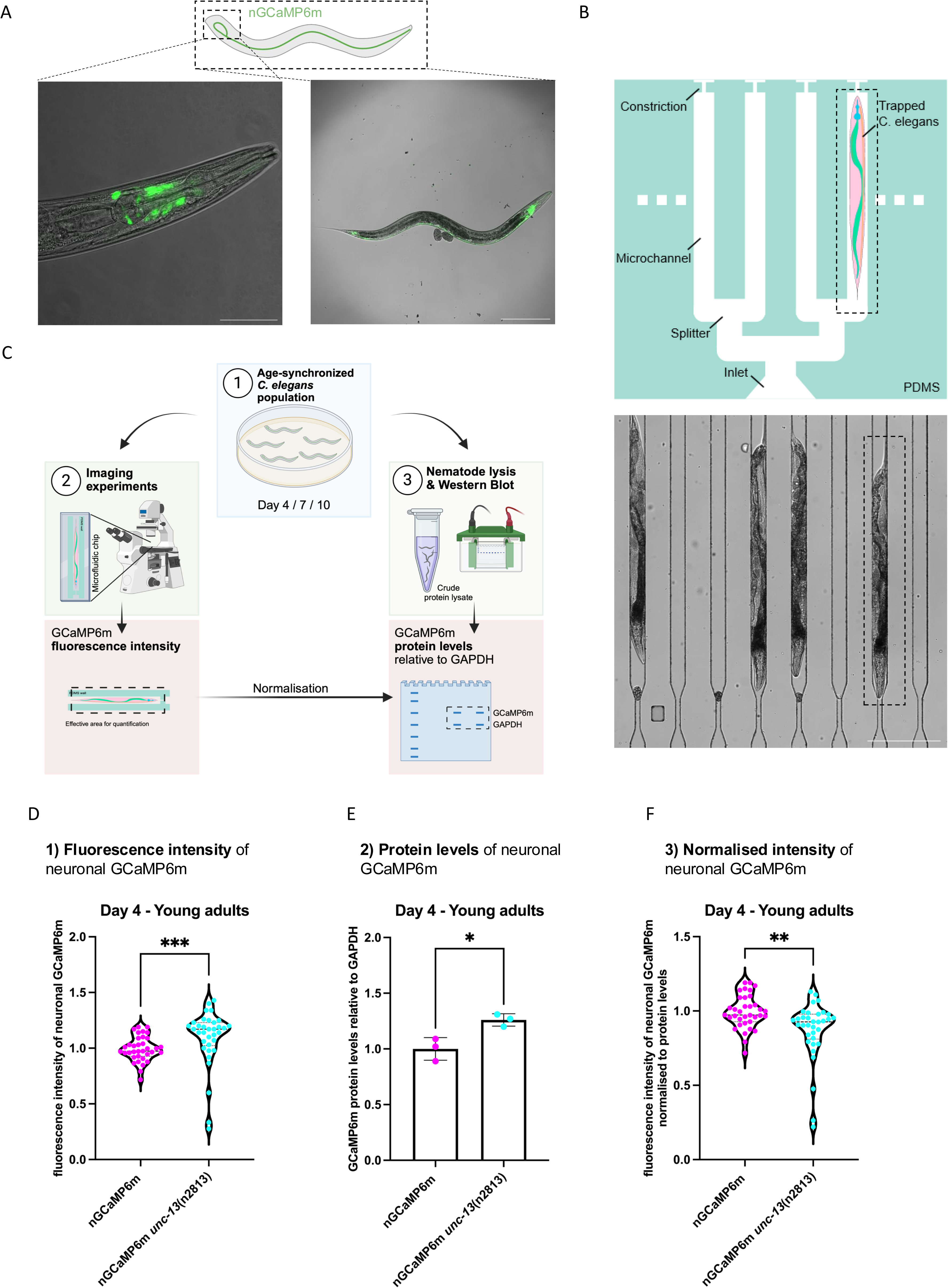
Proof of concept: assessment of neuronal activity by quantifying GCaMP6m intensity levels of living animals in microfluidic devices. A. Representative confocal fluorescent images of a young adult animal (4 days old) of the novel pan-neuronal GCaMP6m strain (nGCaMP6m). The image on the right is 100-fold magnified (scale bar is 200 μm). Inset shows respective close-up of the head region depicted on the left. Close-up image is 400-fold magnified (scale bar is 50 μm). B. Schematic representation of the design of the polydimethylsiloxane (PDMS) microfluidic device used for the imaging of living *C. elegans* (top image). Animals enter the device at the inlet by gravity and are sucked into the channels by applying gentle negative pressure. The constriction prevents animals from escaping at the opposing side. Inside the channels the animals are mechanically trapped along their body and C. *elegans* are straightened. The device allows imaging of multiple animals simultaneously. The image depicted at the bottom is 100-fold magnified with a scale bar of 200 µm. C. Workflow for the quantification of GCaMP6m fluorescence intensity in living *C. elegans* as a readout for the assessment of neuronal activity. Generation of age- synchronized cohorts of young adults (day 4 of life) and older animals (day 7 and 10 of life) (1). Imaging experiments are performed using PDMS microfluidic devices to record GCaMP6m fluorescence intensities from alive animals. In the subsequent analysis step GCaMP6m fluorescence intensities are quantified from single snapshot images. Age-synchronized nematodes are lysed to obtain the crude protein fraction (2). Lysates are analyzed by Western blots. GCaMP6m protein levels are quantified relative to GAPDH and are used to normalize GCaMP6m fluorescence intensities. D. Violine dot plot of the average GCaMP6m fluorescence of young adult animals (day 4-old) of the neuronal GCaMP6m strain (nGCaMP6m) and the nGCaMP6m *unc- 13*(n2813) mutant strain. The mutant *unc*-13(n2813) (strain MT8004) was crossed with the nGCaMP6 strain and GCaMP6m intensities were measured in animals using microfluidic devices with a widefield fluorescence microscope as outlined in Fig. 1B+C. GCaMP6m intensities were quantified using Fiji. Every dot represents the neuronal GCaMP6m fluorescence intensity of a single animal of nGCaMP6m (magenta) and nGCaMP6m *unc-13*(n2813) (turquoise). n = 3 and N = 35 animals. Mann-Whitney test was performed to assess significance (***= p ≤ 0.001). E. Quantification of GCaMP6m proteins levels by Western blot from crude protein lysates of young adult (day 4-old) nGCaMP6m (magenta) and nGCaMP6m *unc- 13*(n2813) (turquoise) animals. Scatter dot plot shows quantification of GCaMP6m protein levels relative to GAPDH from 3 independent cohorts. Unpaired Student’s t- test with Welch’s correction was performed to assess significance (* = p ≤ 0.05). Full- length and annotated Western blots used for the quantification are shown in fig. S6A- C. F. Violine dot plot of the average GCaMP6m fluorescence intensity normalized to the GCaMP6m protein level of young adult animals (day 4-old) of the neuronal GCaMP6m strain (nGCaMP6m) and the nGCaMP6m *unc-13*(n2813) mutant strain. GCaMP6m intensities as depicted in (D) were multiplied with the ratio between nGCaMP6m protein levels of the nGCaMP6m strain and nGCaMP6m unc-13(n2813) mutant strain. Every dot represents the neuronal GCaMP6m fluorescence intensity normalized to GCaMP6m protein levels of a single animal of nGCaMP6m (magenta) and nGCaMP *unc-13*(n2813) (turquoise). n = 3 and N = 35 animals. Mann-Whitney test was performed to assess significance (**= p ≤ 0.01).

The workflow of the analysis using the microfluidic channels is depicted in figure 1C and starts with the synchronization of the nematodes. In this study, we analyzed young adult (4 days after hatching) animals as well as day 7, and day 10 old nematodes to reflect the progression of aging. Notably, animals expressing disease-associated aggregation-prone proteins display a reduced lifespan (median of 9 days for Aβ_1-42_ animals) and the time points of our analysis were chosen to account for the shortened lifespan of the disease models ^11^. For the GCaMP recordings, the nematodes were trapped into the microfluidic channels and their fluorescence intensity was recorded. The nematodes in the channels remain alive and are not harmed during this procedure. The animals can move with their head and tail and still lay eggs, suggesting that the physiological behavior is not impeded. Further, the experiment, starting from the loading of multiple nematodes into the microchannels to imaging the last one at the microscope, takes less than 30 minutes.

To determine the level of calcium-based neuronal signaling in the nematodes, we quantified the fluorescence intensity as well as the protein levels that report on the abundance of the expressed GCaMP protein (Figs. 1C-E and Fig. S2) and then normalized the fluorescence intensity to the GCaMP protein levels (Fig. 1F). This normalization was used for all subsequent analyses in this study to account for variations in GCaMP expression levels at different ages of the animals and different strains. To further verify our observations above, we also performed this analysis in *unc-13* mutant animals that are impaired in synaptic vesicle fusion ^14^ and as expected, they showed a significant decrease in the GCaMP intensity (Fig. 1F). As additional control, we applied the calcium channel inhibitor Nemadipine A to GCaMP-expressing control nematodes ^15^. Nemadipine A treated animals exhibited a significant decrease of the GCaMP fluorescence (Fig. S3). We conclude that our experimental setup can be used to report on the average neuronal activity of living *C. elegans*.

### Neuronal function declines prior to the onset of significant Aβ_1-42_ aggregation

Aβ_1-42_-expressing animals show an age-dependent increase in Aβ_1-42_ aggregation, propagation and severe proteotoxicity ^11^. To correlate the expression and aggregation of Aβ_1-42_ with its effect on neuronal function, we crossed the strain with the pan-neuronal GCaMP (nGCaMP6m) reporter with the strain expressing neuronal Aβ_1-42_ (nAβ) or control animals that only express the fluorescent protein mScarlet in the neurons (termed nmScarlet) (Fig. 2A). We then measured the neuronal activity at two time points, *i.e.,* on day 4 of life (young adults) with the start of Aβ_1-42_ aggregation and on day 7 of life (older adults) when the animals show severe Aβ_1-42_ aggregation as demonstrated previously ^11^ (Fig. 2B, Fig. S4A and Fig. S5A,C+D). Notably, compared to the control, we observed a severe reduction in neuronal activity already in day 4 old Aβ_1-42_ animals. Day 7 old Aβ_1-42_ animals showed, as expected, a reduction in neuronal activity. At this age, Aβ_1-42_ aggregation affects all neurons, resulting in severe proteotoxicity as shown previously ^11^. Surprisingly, the neuronal activity on day 4 was lower compared to that of day 7 for Aβ_1-42_ animals (Fig. 2B). This may seem contradictory at first glance, but could be explained with a survivorship bias as the median lifespan of Aβ_1-42_ animals is 9 days ^11^. Thus, animals that are more severely affected by the aggregation and proteotoxicity of Aβ_1-42_ may have already died before reaching day 7.

**Figure 2.**
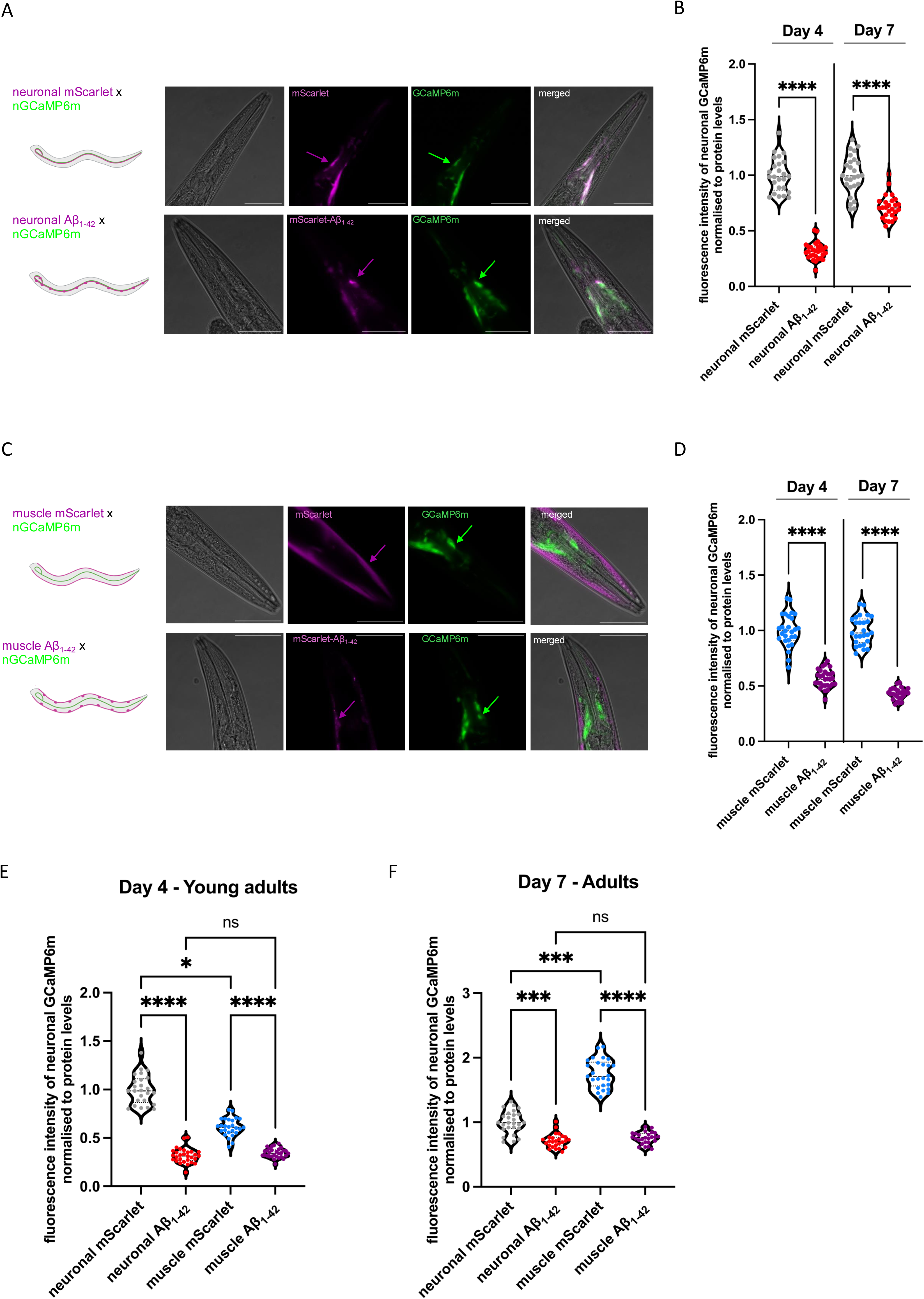
Decrease of neuronal activity in Alzheimer’s disease models precedes the accumulation of Aβ_1-42_ aggregates. A. Representative confocal fluorescent images of a young adult animal (day 4-old) of the nmScarlet x nGCaMP6m cross (top panel) and of the cross nAβ_1-42_ x nGCaMP6m (bottom panel). Shown are close-up images of the head region of the individual channels and of the merge. Images are 400-fold magnified and scale bars are 50 μm. B. Violine dot plot of the average GCaMP6m fluorescence intensity normalized to the GCaMP6m protein level of young adult animals (day 4-old) and adult animals (day 7-old) of the control strain expressing neuronal mScarlet (nmScarlet) and nAβ_1-42_. GCaMP6m intensities as depicted in figure S4A were multiplied with the ratio between nGCaMP6m protein levels of the nmScarlet x nGCaMP6m cross and nAβ_1-42_ x nGCaMP6m cross for Day 4 and 7 respectively. GCaMP6m protein quantification is shown in figure S5A and the corresponding full-length Western blots in figure S5C+D. Every dot represents the neuronal GCaMP6m fluorescence intensity normalized to GCaMP6m protein levels of a single animal of nmScarlet (grey) and nAβ_1-42_ (red). n = 3 and N = 25-29 animals. Significance was assessed between nmScarlet Day 4 and nAβ_1-42_ Day 4 and nmScarlet Day 7 and nAβ_1-42_ Day 7, respectively by unpaired Student’s t-test with Welch’s correction (**** = p ≤ 0.0001). C. Representative confocal fluorescent images of a young adult animal (day 4-old) of the mmScarlet x nGCaMP6m cross (top panel) and of the mAβ1-42 x nGCaMP6m cross (bottom panel). The mAβ_1-42_ strain as well as the respective mmScarlet control strain were crossed with the nGCaMP6 strain. Shown are close-up image of the head region of the individual channels and of the merge. Images are 400-fold magnified and scale bars are 50 μm. D. Violine dot plot of the average GCaMP6m fluorescence intensity normalized to the GCaMP6m protein level of young adult animals (day 4-old) and adult animals (day 7- old) of the control strain expressing muscle mScarlet (mmScarlet) and mAβ_1-42_. GCaMP6m intensities were measured in alive animals using microfluidic devices with a widefield fluorescence microscope. GCaMP6m intensities as depicted in figure S4B were multiplied with the ratio between nGCaMP6m protein levels of the mmScarlet x nGCaMP6m cross and mAβ_1-42_ x nGCaMP6m cross for Day 4 and 7 respectively. GCaMP6m protein quantification is shown in figure S5B and the corresponding full- length Western blots in figure S5C+D. Every dot represents the neuronal GCaMP6m fluorescence intensity normalized to GCaMP6m protein levels of a single animal of mmScarlet (turquoise) and mAβ_1-42_ (purple). n = 3-4 and N = 26-30 animals. Significance was assessed between mmScarlet Day 4 and mAβ_1-42_ Day 4 and mmScarlet Day 7 and mAβ_1-42_ Day 7 respectively by unpaired Student’s t-test with Welch’s correction (**** = p ≤ 0.0001). E. Violine dot plot of the average GCaMP6m fluorescence intensity normalized to the GCaMP6m protein level of young adult animals (day 4-old) of the nmScarlet, nAβ_1- 42_, mmScarlet and mAβ_1-42_ strain. GCaMP6m intensities were measured in alive animals using microfluidic devices with a widefield fluorescence microscope and were multiplied with the ratio between nGCaMP6m protein levels of the nmScarlet x nGCaMP6m cross and nAβ_1-42_ x nGCaMP6m / mmScarlet x nGCaMP6m / mAβ_1-42_ x nGCaMP6m, respectively to allow comparison between tissues. Every dot represents the neuronal GCaMP6m fluorescence intensity normalized to GCaMP6m protein levels of a single animal of nmScarlet (grey), and nAβ_1-42_ (red), mmScarlet (turquoise) and mAβ_1-42_ (purple). n = 3-4 and N = 25-30 animals. The graph shows the same data sets depicted in individual graphs in B and D but compares the aggregation of Tau in the different strains of the same age. Significance was assessed by Kruskal-Wallis test with Dunn post hoc test (**** = p ≤ 0.0001). F. Violine dot plot of the average GCaMP6m fluorescence intensity normalized to the GCaMP6m protein level of adult animals (day 7-old) of the nmScarlet, nAβ_1-42_, mmScarlet and mAβ_1-42_ strain. GCaMP6m intensities were measured in alive animals using microfluidic devices with a widefield fluorescence microscope and were multiplied with the ratio between nGCaMP6m protein levels of the nmScarlet x nGCaMP6m cross and nAβ_1-42_ x nGCaMP6m / mmScarlet x nGCaMP6m / mAβ_1-42_ x nGCaMP6m, respectively to allow comparison between tissues. Every dot represents the neuronal GCaMP6m fluorescence intensity normalized to GCaMP6m protein levels of a single animal of nmScarlet (grey), and nAβ_1-42_ (red), mmScarlet (turquoise) and mAβ_1-42_ (purple). n = 3 and N = 26 animals. The graph shows the same data sets depicted in individual graphs in B and D but compares the aggregation of Tau in the different strains of the same age. Significance was assessed by Kruskal-Wallis test with Dunn post hoc test (**** = p ≤ 0.0001).

Next, we asked the question how Aβ_1-42_ expressed in a distal tissue affects neuronal activity. To address this, we made use of a muscle Aβ_1-42_ strain that expresses the Aβ_1-42_ peptide in body wall muscles (mAβ) ^11^ and crossed this strain as well as muscle mScarlet control nematodes (termed mmScarlet) with animals expressing neuronal GCaMP (Fig. 2C). Surprisingly, we observed a similar reduction in neuronal activity when Aβ_1-42_ is expressed in muscle tissue, which also already occured in young adult animals (day 4). Upon closer inspection, we noticed that the detrimental effect of Aβ_1-42_ on neuronal activity is on day 7 more pronounced when Aβ_1-42_ is expressed in muscle compared to a neuronal expression (Figs. 2D-F, Fig. S4B and Fig. S5B-D). This could be a direct effect of propagated Aβ_1-42_ from muscle to neurons and / or transduced by a signaling cascade as aging progresses ^16^. We conclude from this data set that the neuronal activity decreases early in Aβ_1-42_-expressing animals and occurs before a significant accumulation of Aβ_1-42_ aggregates. In addition, the neuronal activity seems equally affected by distally expressed Aβ_1-42_.

### Generation of neuronal *C. elegans* Tau models based on the human Tau-0N4R isoform

The obtained data with the Aβ_1-42_ model posed the question whether the early decline of neuronal activity in AD animals is specific for Aβ_1-42_ or a general phenomenon in response to aggregating disease-associated amyloid proteins. We set out to perform the same analysis with the second amyloid protein that is associated with AD, Tau. Tau is a microtubule-binding, intracellular protein that can become hyperphosphorylated and can form neurofibrillary tangles ^17^. Several point mutations have been identified in familial cases of AD that render Tau aggregation prone ^18^. We generated strains that express the human 0N4R variant Tau in a non-modified form that is referred to as wild-type Tau (Tau^WT^) and a mutant Tau form harboring two patient-derived mutations, P301L and V337M (Tau^P301L,V337M^) in the neurons (Fig. 3A). Analogously to the Aβ_1-42_ model, Tau is expressed in an untagged and sub-stoichiometrically mScarlet-tagged manner using an IRES element for its synthesis (Figs. 3B+D + Fig. S6) ^11^. The sub-stoichiometric tagging with mScarlet limits a perturbation of the amyloid fibril formation by the fluorescent protein yet enables a visualization and assessment of Tau aggregation by fluorescence lifetime imaging microscopy (FLIM) in a non-invasive manner. Surprisingly, neuronal Tau expression increased with the progression of aging from day 4 (young adult animals) up to day 10 (past their fertile period) for both Tau^WT^ and Tau^P301L,V337M^ (Figs. 3C+E + Fig S7A) and differed in that regard from nAβ animals that showed decreased Aβ_1-42_ expression with aging (Figs. S7B+C). The increased abundance of Tau with aging is likely due to an increased stability of the protein as the transcription increased only for Tau^P301L,V337M^ between days 4 and 7 and did not further increase to day 10 and was not changed at all for Tau^WT^ (Fig. 3F).

**Figure 3.**
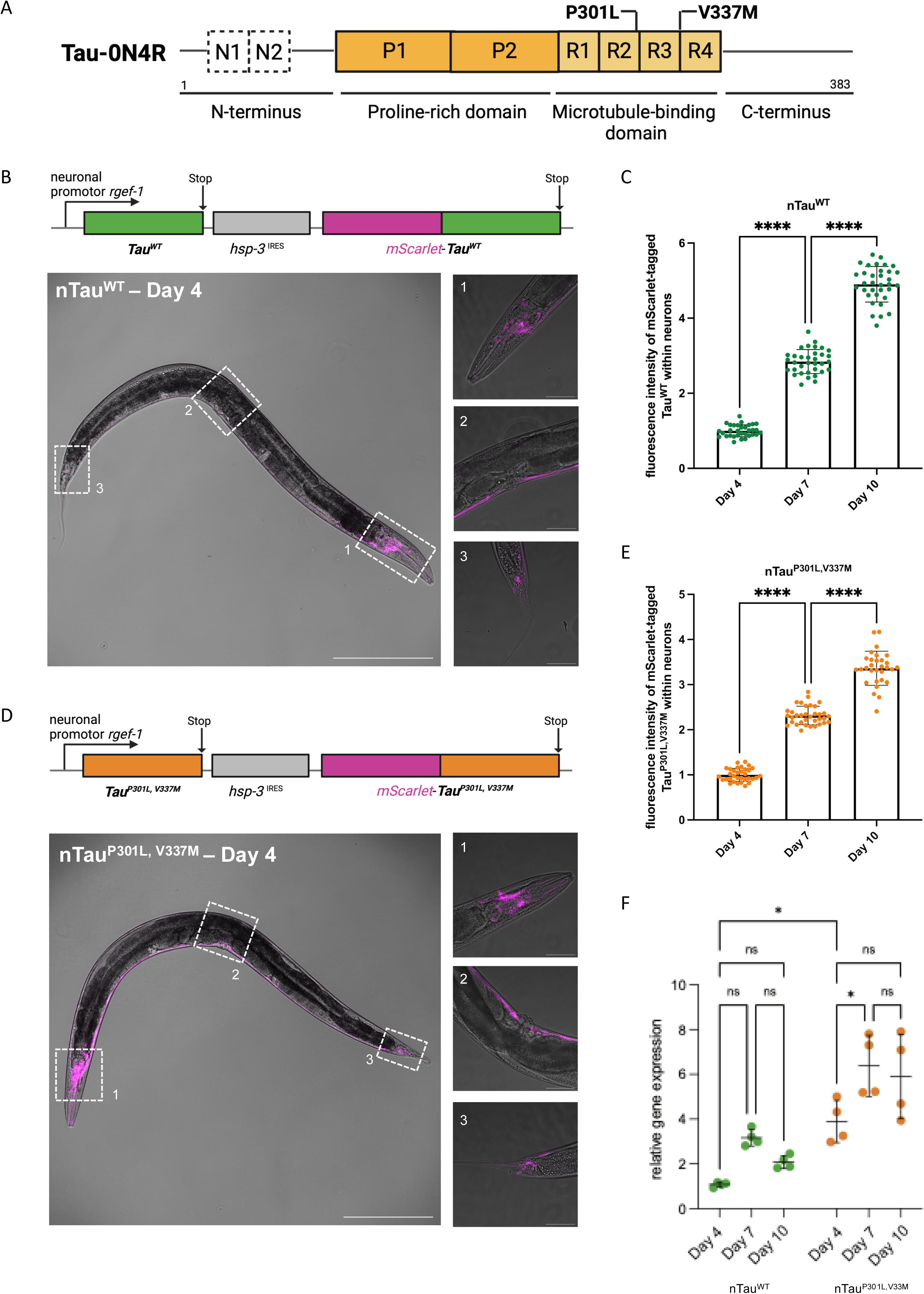
New *C. elegans* Tau model. A. Protein domain structure of the human Tau protein isoform 0N4R used for over- expression. The N-terminus of Tau-0N4R lacks the two N-terminal domains N1 and N2 (dashed squares) followed by a proline-rich domain, the four repeat domains (R1- R4) of the microtubule-binding domain and the C-terminus. The position of the two point-mutations Proline-to-Leucine at residue 301 (P301L) and Valine-to-Methionine at residue 337 (V337M) that are associated with frontotemporal dementia (FTD) are indicated and used to generate the mutant Tau strain. B. Schematic representation of the neuronal Tau^WT^ (nTau^WT^) expression construct and corresponding confocal fluorescent images of a young adult (day 4-old) animal. Untagged Tau^WT^ and mScarlet-tagged Tau^WT^ are pan-neuronally expressed driven by the *rgef-1* promotor. Cap-dependent translation leads to an overexpression of untagged Tau^WT^ and *hsp-3*^IRES^ cap-independent translation allows for sub- stoichiometric expression of mScarlet-Tau^WT^. Image on the left is 100-fold magnified (scale bar is 200 μm). Insets 1-3 show the respective close-ups of the head-, mid body and tail region. Close-up images on the right are 400-fold magnified (scale bars are 50 μm). C. Quantification of mScarlet-Tau^WT^ fluorescence intensity with the progression of aging of day 4-, 7- and 10-old nematodes of nTau^WT^ strain. Confocal fluorescent images of three cohorts of total 32-35 nematodes were recorded, and fluorescence intensities were quantified by Fiji. Data are displayed as mean fluorescence intensity ± SD. Significance was tested by one-way ANOVA + Bonferroni post hoc test (**** = p < 0.0001). D. Schematic representation of the neuronal Tau^P301L,V337M^ (nTau^P301L,V337M)^ construct and corresponding fluorescent confocal microscopy images of a young adult (day 4-old) animal. The presented operon structure follows the same principle of expression as described in B. Image on the left is 100-fold magnified (scale bar is 200 μm). Insets 1-3 show the respective close-ups of the head-, mid body and tail region. Close-up images on the right are 400-fold magnified (scale bars are 50 μm). E. Quantification of mScarlet-Tau^P301L,V337M^ fluorescence intensity with the progression of aging of day 4-, 7- and 10-old nematodes of nTau^P301L, V337M^ strain. Confocal fluorescent images of three cohorts of total 32-35 nematodes were recorded, and fluorescence intensities were quantified by Fiji. Data are displayed as mean fluorescence intensity ± SD. Significance was tested by one-way ANOVA + Bonferroni post hoc test (**** = p < 0.0001). F. qRT-PCR analysis of Tau expression. The graph depicts the relative human Tau expression of JKM150 (Tau^WT^) and JKM151 (Tau^P301L,V337M^) on day 4, day 7 and day 10 of life. The expression data were normalized to the average expression of three reference genes (*act-1*, *lmn-1* and *eif-3C*). Three independent cohorts of approximately were analyzed. Shown is the mean ± SD. Ratio-paired t test was performed. n.s. = non-significant; * = p ≤ 0.05

### Mutant Tau shows age-dependent aggregation and propagation

Next, we analyzed the aggregation of Tau using FLIM (Fig. 4A). FLIM is an advanced microscopy technique capable of providing a deeper understanding of the molecular environment of a fluorophore. Amyloid formation leads to the quenching of the mScarlet fluorophores that are fused to Tau, which reduces the fluorophore’s fluorescence lifetime (τ) ^19^. Thus, the fluorescence lifetime can be used as readout for the aggregation of Tau ^20^. The control (nmScarlet) and Tau^WT^ were soluble whereas Tau^P301L,V337M^ aggregated already in young adult animals (Figs. 4B+C). The aggregation of Tau^P301L,V337M^ is also depicted by the color change in the false-colored images where blue areas indicate regions of low lifetime (= aggregates) and reflected by the overall decreased τ values of the head neurons (Fig. 4A). We observed further that the aggregation of mutant Tau (Tau^P301L,V337M^) increased with age while the control and Tau^WT^ remained soluble (Fig. 4C). These results highlight that the increased abundance that was observed for both Tau^WT^ and Tau^P301L,V337M^ with aging (Figs. 3C+E) is not necessarily leading to a higher aggregation propensity and that instead the P301L and V337M mutations cause Tau aggregation.

**Figure 4.**
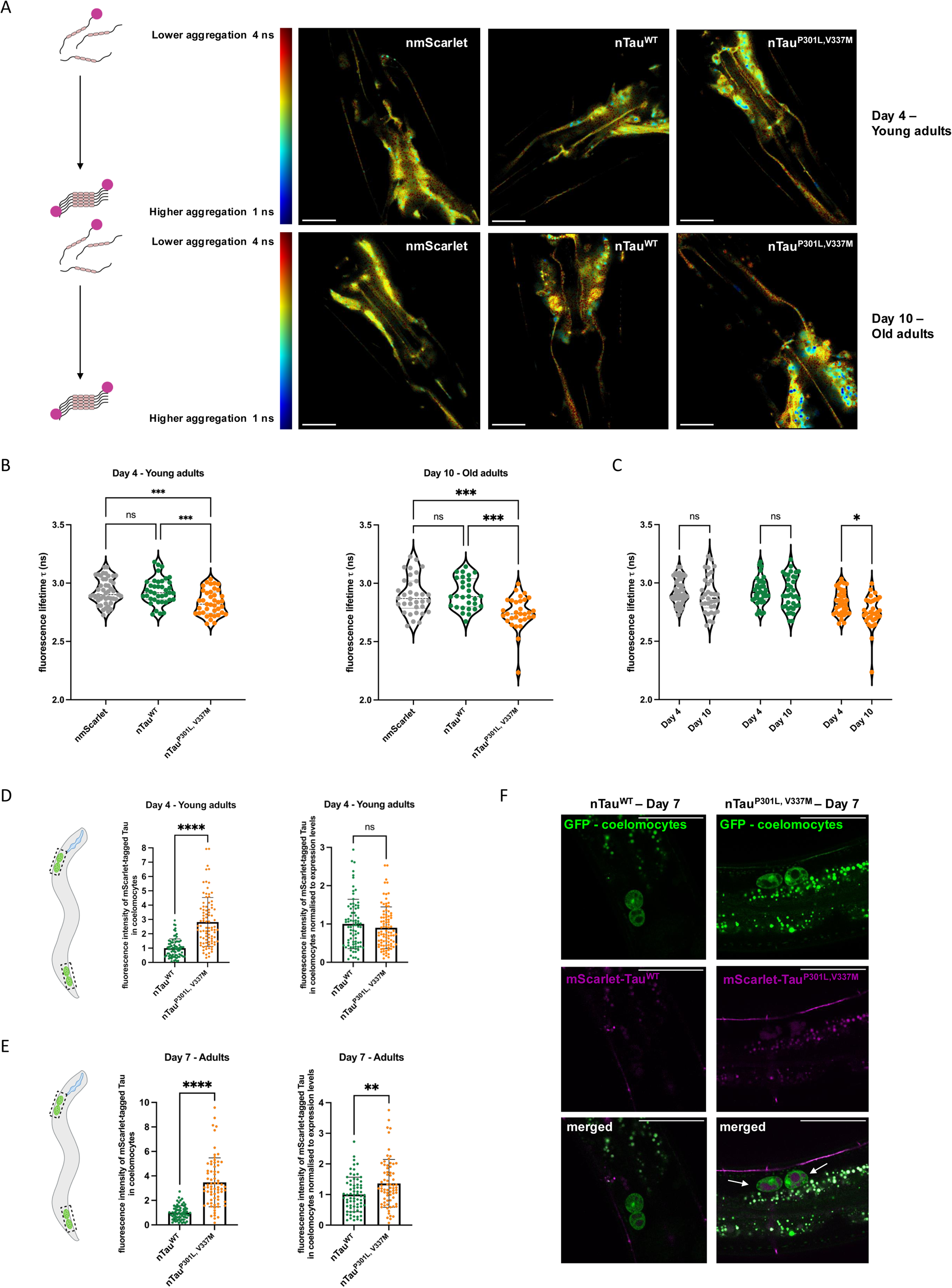
Mutant Tau shows age-dependent aggregation and propagation. A. Representative TCSPC-FLIM images of nmScarlet, nTau^WT^ and nTau^P301L,V337M^. Head neurons of young adult (day 4-old) animals (upper row) and old adult (day 10- old) animals (lower row) were analyzed. Images depict the pixel-wise fluorescent lifetime (Δ) merged with the fluorescent intensity. Fluorescence lifetime is false- colored: blue represents low fluorescent lifetimes (1 ns) showing regions with aggregated Tau and red represents high fluorescent lifetimes (4 ns) showing region with soluble Tau species that are schematically depicted on the left. Scale bars are 20 μm. B. Violine dot plot of the average fluorescent lifetime (Δ) of young adult animals (left) and old adult animals (right) of nmScarlet, nTau^WT^ and nTau^P301L,V337M^. Data displays average fluorescent lifetimes ± SD of nmScarlet (grey), nTau^WT^ (green) and nTau^P301L, V337M^ (orange). Every dot represents the average fluorescent lifetime for the head neurons of one single nematode. Three independent cohorts of in total 31-47 nematodes were analyzed. Significance was tested by one-way ANOVA + Bonferroni post hoc test for young adult animals and by Kruskal-Walls test with Dunn post hoc test for old animals (ns = p > 0.05; *** = p ≤ 0.001). C. Violine dot plot of the average fluorescent lifetime (Δ) of young adult animals (day 4-old) and old adult animals (day-10 old) of nmScarlet, nTau^WT^ and nTau^P301L,V337M^. The graph shows the combined data sets depicted in individual graphs of (B). Data displays average fluorescent lifetimes ± SD of nmScarlet (grey), nTau^WT^ (green) and nTau^P301L,V337M^ (orange). Every dot represents the average fluorescent lifetime for the head neurons of one single nematode. Three independent cohorts of in total 31-47 animals were analyzed. Significance was tested by two-way ANOVA + Bonferroni post hoc test (ns = p > 0.05; * = p ≤ 0.05). D. Scatter dot plot of the average Tau levels in the coelomocytes of nTau^WT^ and nTau^P301L,V337M^ young adult animals (left graph) as well as normalized to the Tau expression levels (right graph). For identification of coelomocytes nTau^WT^ and nTau^P301L,V337M^ were crossed with ZIM1048 strain expressing GFP in coelomocytes. We excluded day 10 analyses due to the increased autofluorescence at that age. Tau fluorescence levels were quantified in the head and the tail coelomocytes by Fiji analysis of confocal images. Every dot represents the level of Tau in head or the tail coelomocytes for nTau^WT^ (green) and nTau^P301L, V337M^ (orange). Fluorescence intensities were normalized to day 4 of the signals of nTau^WT^. Three cohorts of in total 32-38 nematodes were analyzed. Significance was assessed by Mann-Whitney test (ns = p > 0.05; **** = p ≤ 0.0001). E. Scatter dot plot of the average Tau levels in the coelomocytes of nTau^WT^ and nTau^P301L,V337M^ adult animals (left graph) as well as normalized to the Tau expression levels (right graph). Tau fluorescence levels were quantified in the head and the tail coelomocytes by Fiji analysis of confocal images. Every dot represents the level of Tau in head or the tail coelomocytes for nTau^WT^ (green) and nTau^P301L,V337M^ (orange). Fluorescence intensities were normalized to day 7 of the signals of nTau^WT^. Three cohorts of in total 37-38 nematodes were analyzed. Significance was assessed by Mann-Whitney test (** = p ≤ 0.01; **** = p ≤ 0.0001), F. Confocal fluorescent images of nTau^WT^ (left) and nTau^P301L,V337M^ (right) adult animals (day 7-old) crossed with the coelomocyte marker (*unc-122p::gfp*) strain ZIM1048. Scale bars are 50 μm. Arrows in the merged panel of Tau^P301L,V337M^ point to coelomocytes that show incorporated magenta, fluorescent material representing mScarlet-tagged Tau^P301L,V337M^.

A hallmark of neurodegenerative diseases is the spreading of the disease-associated amyloid proteins. We hence analyzed spreading of Tau^P301L,V337M^ to other tissues. To quantify this, we measured the Tau levels in the coelomocytes, scavenger cells that take up extracellular material from the body cavity. Of note, only proteins that are released from the neurons as part of the spreading and propagation of Tau can enter the extracellular space and get endocytosed by coelomocytes where the proteins can be subjected to lysosomal degradation. We crossed the Tau strains with a strain that expresses GFP in the coelomocytes that serves as a marker to easily identify them and then quantify Tau levels in the coelomocytes. On day 4 and day 7, we detected a 2.8-fold (day 4) or even 3.5-fold (day 7) increase of mutant Tau compared to Tau^WT^ in the coelomocytes (Figs. 4D-F). When we normalized the abundance in the coelomocytes to the Tau expression levels, it became clear that the spreading of Tau^P301L,V337M^ was still significantly higher than that of Tau^WT^ yet only in older animals (Fig. 4D+E right plot), when we also observed strong aggregation of Tau^P301L,V337M^ by FLIM (Figs. 4A-C).

### Aggregation of Tau is associated with severe organismal defects

Next, we performed a phenotypic analysis of the new Tau strains to comprehensively characterize them and to assess the proteotoxicity of Tau. Tau^P301L,V337M^ expression led to a significantly reduced lifespan with a median lifespan of 14.2 days in comparison to the median lifespan of nmScarlet control of 16.5 days (Fig. 5A, Fig. S10). Further, we assessed several physiological parameters that report on the fitness of animals. We observed a severe 48% reduction in the number of progenies, from a mean number of 267 offspring for nmScarlet control animals to 139 for the Tau^P301L,V337M^ animals (Fig. 5B). Additionally, the development of the mutant Tau animals was severely delayed. This was already apparent on day 3 of life and even more on day 4 of life, when about 48 % of all mutant Tau animals were still in a larval stage while the control animals all reached adulthood (Fig. 5C). Alongside this, a chemotaxis assay revealed that mutant Tau animals were impaired to respond to the volatile attractant 2% Benzaldehyde with a mean chemotaxis index of 0.76 as compared to N2 wild-type and Tau^WT^ animals showing a mean chemotaxis index of 0.89 and 0.9 respectively (Fig. 5D). Finally, motility, as assessed by thrashing frequency in liquid media, was reduced in older mutant Tau animals with a mean thrashing frequency of 1.5 Hz compared to nmScarlet control animals with a thrashing frequency of 2 Hz (Fig. 5E). Notably, we also observed toxicity for the expression of Tau^WT^, which was unexpected as Tau^WT^ does not aggregate (Figs. 4A+B). We noticed that Tau^WT^ elicited its detrimental effect only in assays that report on longer periods of the nematodes’ life e.g. lifespan (up to day 25) and fecundity assay (up to day 8), whereas we did not observe defects in development on days 3 and 4 of life (Fig. 5C), chemotaxis on day 4 (Fig. 5D) and motility on day 4 (Fig. 5E). Defects in motility could only be detected on day 10 (Fig. 5E). Thus, the expression of Tau^WT^ exerted a negative and aggregation-independent effect on the physiology of adult nematodes (Figs. 4A- C).

**Figure 5.**
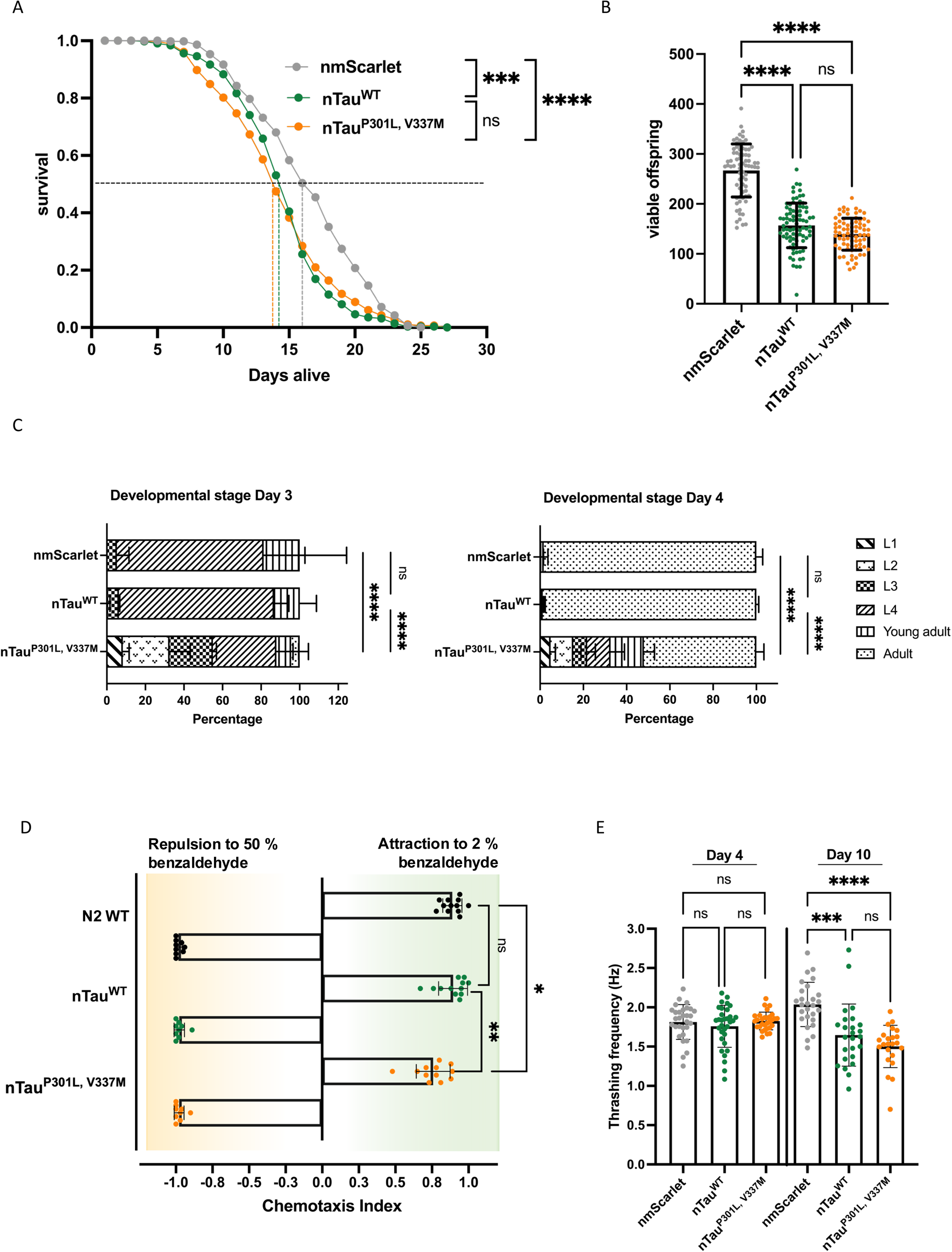
Aggregation of Tau is associated with severe organismal defects. A. Assessment of the lifespan of nmScarlet, nTau^WT^ and nTau^P301L,V337M^ animals. Graph shows the cumulative survival probability (survival) versus age (days alive) of nmScarlet (grey), nTau^WT^ (green) and nTau^P301L,V337M^ (orange). Three independent cohorts of 150-180 nematodes each were analyzed, and significance was tested by log-rank test using Oasis2 online tool (ns = p > 0.05; *** = p ≤ 0.001, **** = p ≤ 0.0001). B. Fecundity analysis of nmScarlet, nTau^WT^ and nTau^P301L,V337M^ animals. Scatter dot plot displays average number of viable offspring of nmScarlet (grey), nTau^WT^ (green) and nTau^P301L,V337M^ (orange). Every dot represents the absolute number of viable offspring of a single nematode. Significance was assessed by one-way ANOVA + Bonferroni post hoc test (ns = p > 0.05; **** = p ≤ 0.0001). Three independent cohorts of in total 74-83 nematodes were tested. C. Developmental assay of nmScarlet, nTau^WT^ and nTau^P301L,V337M^ animals. Graph displays percentage of developmental stages of the nematodes within a population at day 3 (left) and day 4 (right) of life. Three independent cohorts of 40-60 nematodes each were analyzed. Kruskal-Wallis test was employed to test significance between fractions of L4 animals (left) and adult animals (right) (**** = p ≤ 0.0001). D. Assessment of chemotaxis towards 50% (repulsion) and 2% (attraction) benzaldehyde of nTau^WT^ and nTau^P301L,V337M^ animals. Scatter plot displays chemotaxis index of nTau^WT^ (green) and nTau^P301L,V337M^ (orange) whereas each dot represents the calculated chemotaxis index for one technical replicate. 3-4 independent cohorts of each 200-300 nematodes (young adults) were tested, divided into three technical replicates. Significance was assessed by Kruskal-Wallis test with Dunn post hoc test (ns = p > 0.05; * = p ≤ 0.05; **= p ≤ 0.01). E. Scatter dot plot of the thrashing capability of nmScarlet (grey), nTau^WT^ (green) and nTau^P301L,V337M^ (orange) animals at day 4 (left) and day 10 (right) of life. Every dot represents the trashing frequency of a single nematode. Three cohorts of 8-15 nematodes per strain were recorded for 20 s and thrashing frequency was analyzed from the recorded videos for each animal individually. Significance for day 4 and 10 respectively was assessed by Kruskal-Wallis test with Dunn post hoc test (ns = p > 0.05; *** = p ≤ 0.001; **** = p ≤ 0.0001).

We set out to further study the neuronal ectopic expression of human Tau^WT^ in *C. elegans*. The endogenous *C. elegans* Tau ortholog, PTL-1, is expressed in several neurons, most strongly in ALM, AVM, and PLM touch neurons, and is required for neuronal integrity ^21,22^. PTL-1A and PTL-1B are two isoforms and show less than 50% sequence identity with human Tau (44% for PTL-1A and 47% for PTL-1B) that is however restricted to the microtubule binding domains (Fig. S8B) ^23^. Mutation of *ptl-1* leads to neuronal morphological defects and a shortened lifespan ^21^. We could reproduce these data and observed a reduction in the lifespan of *ptl-1* mutant (*ok621*; Fig. S8A). It has previously been shown that the shortened lifespan of *ptl-1* mutant animals can be rescued by ectopic expression of *ptl-1* ^21^. However, when we crossed the *ptl-1* mutant with our newly generated neuronal human Tau^WT^ strain, we observed an even further reduction in lifespan from a median lifespan of 14.45 days (*ptl-1* deletion mutant) and 14.33 (pan-neuronal overexpression human Tau^WT^) to 11.23 days for the cross of *ptl-1* mutant and neuronal human Tau^WT^ (Fig. S8A). These data show that human Tau differs from its *C. elegans* homolog and cannot substitute for it. Thus, the cross of *ptl-1* mutant and human Tau^WT^ is deficient for the endogenous Tau protein and expresses a protein that is dominantly negative on the physiology of the nematode. Importantly, Tau^WT^ does not exert its toxicity by aggregation (Fig. 4A+B). We thus argue that Tau^WT^ is not a suitable control for the aggregation-associated toxicity of Tau^P301L,V337M^ and instead the mScarlet-expressing strain should be used as control for the toxic gain-of-function of amyloid fibril forming Tau^P301L,V337M^. Tau^P301L,V337M^ on the other hand showed in all behavioral assays a proteotoxic effect. The defects in neuronal function as reported in the chemotaxis assay where Tau^P301L,V337M^ animals were impaired to respond to an attractant, led us to hypothesize a reduced neuronal activity in the subsequent GCaMP analyses.

### Neuronal function declines prior to the onset of significant Tau aggregation

To correlate aggregation of Tau^P301L,V337M^ with neuronal activity, we crossed the Tau lines with the GCaMP reporter and quantified the GCaMP fluorescence analogous to the analyses done for Aβ_1-42_ (Figs. 6A-D, Fig. S4C-E, Fig. S9). Of note, GCaMP expression does not affect the aggregation propensity of Tau^P301L,V337M^ or the lack thereof of mScarlet and Tau^WT^ expressing animals (Fig. S4F). We observed a severe reduction of the relative GCaMP intensity in Tau^P301L,V337M^ already in young adult animals (day 4 of life) that persisted throughout aging (days 7 and 10 of life; Figs. 6B- D). We conclude, the expression of the highly aggregation-prone Tau^P301L,V337M^ results in a severe reduction of neuronal activity already starting in young adult animals similarly to Aβ_1-42_ expression. Importantly, the severe defects in neuronal activity occur early (young adult, day 4 old animals) and precede the continued buildup of Tau^P301L,V337M^ aggregates with aging (Figs. 4A-C). Notably, we also observed reduced GCaMP intensities upon expression of Tau^WT^ in young adult animals (day 4) that however increased with aging (Figs. 6B-D), further supporting that ectopic expression of Tau^WT^ expression alters the physiology of the animal. It could be that calcium signaling and or storage are perturbed in older Tau^WT^-expressing animals. We then asked the question whether the loss of neuronal activity is associated with neurodegeneration. To monitor neuronal integrity, we crossed animals expressing Tau^WT^ and Tau^P301L,V337M^ with a strain expressing GFP in the mechanosensory neurons (*mec-4::gfp*) to visualize neuronal morphology (Fig. 6E). While we did not observe morphological aberrations in Tau^WT^ expressing animals, we did detect swellings and protrusions of neurons in animals expressing Tau^P301L,V337M^ (Fig. 6E). This neurodegeneration is in line with the observed loss of neuronal activity and function of Tau^P301L,V337M^ animals (Figs. 5D + 6A-D).

**Figure 6.**
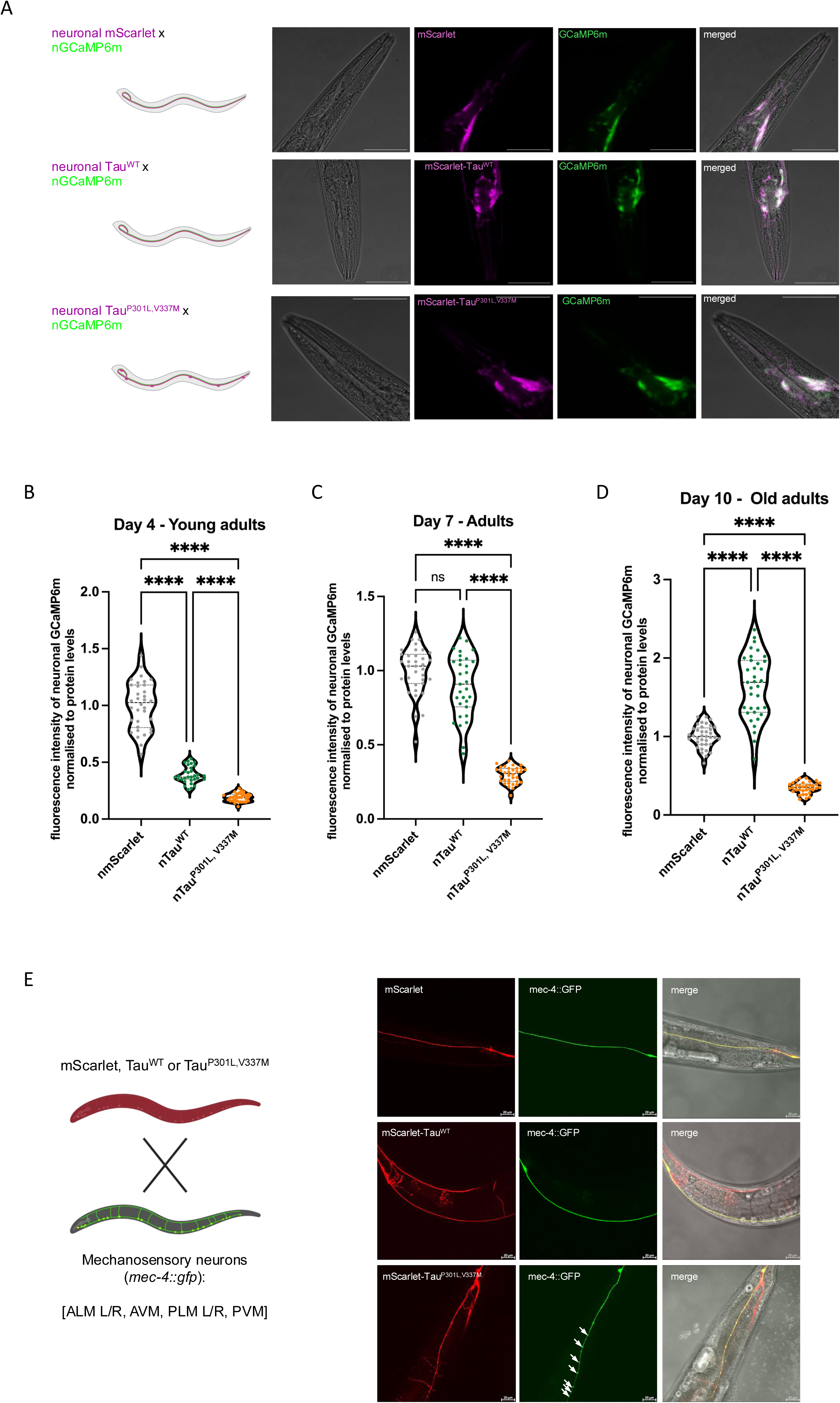
Loss of neuronal function precedes Tau aggregation. A. Representative confocal fluorescent images of a young adult animal (day 4-old) of the nmScarlet x nGCaMP6m, nTau^WT^ x nGCaMP6m, and nTau^P301L,V337M^ x nGCaMP6m cross from left to right. The Tau-expressing strains as well as the nmScarlet control strain were crossed with the nGCaMP6 strain. Shown are close-up image of the head region. Images are 400-fold magnified and scale bars are 50 μm. B. Violine dot plot of the average GCaMP6m fluorescence intensity normalized to the GCaMP6m protein level of young adult animals (day 4-old) of the control strain nmScarlet, nTau^WT^ and nTau^P301L,V337M^. GCaMP6m intensities were measured in alive animals using microfluidic devices with a widefield fluorescence microscope. GCaMP6m intensities as depicted in figure S4C were multiplied with the ratio between nGCaMP6m protein levels of the nmScarlet x nGCaMP6m cross and nTau^WT^ x nGCaMP6m cross / nTau^P301L,V337M^ x nGCaMP6m respectively. GCaMP6m protein quantification is shown in figure S9A and the corresponding full-length Western blots in figure S9D. Every dot represents the neuronal GCaMP6m fluorescence intensity normalized to GCaMP6m protein levels of a single animal of nmScarlet (grey) and nTau^WT^ (green) and nTau^P301L,V337M^ (orange). n = 5 and N = 31-34 animals. Significance was assessed by one-way ANOVA + Bonferroni post hoc test (**** = p ≤ 0.0001). C. Violine dot plot of the average GCaMP6m fluorescence intensity normalized to the GCaMP6m protein level of adult animals (day 7-old) of the control strain nmScarlet, nTau^WT^ and nTau^P301L,V337M^. GCaMP6m intensities were measured in alive animals using microfluidic devices with a widefield fluorescence microscope. GCaMP6m intensities as depicted in figure S4D were multiplied with the ratio between nGCaMP6m protein levels of the nmScarlet x nGCaMP6m cross and nTau^WT^ x nGCaMP6m cross / nTau^P301L,V337M^ x nGCaMP6m respectively. GCaMP6m protein quantification is shown in figure S9B and the corresponding full-length Western blots in figure S9E. Every dot represents the neuronal GCaMP6m fluorescence intensity normalized to GCaMP6m protein levels of a single animal of nmScarlet (grey) and nTau^WT^ (green) and nTau^P301L,V337M^ (orange). n = 4-5 and N = 31-35 animals. Significance was assessed by Kruskal-Wallis test with Dunn post hoc test (ns = p > 0.05; **** = p ≤ 0.0001). D. Violine dot plot of the average GCaMP6m fluorescence intensity normalized to the GCaMP6m protein level of old adult animals (day 10-old) of the control strain nmScarlet, nTau^WT^ and nTau^P301L,V337M^. GCaMP6m intensities were measured in alive animals using microfluidic devices with a widefield fluorescence microscope. GCaMP6m intensities as depicted in figure S4E were multiplied with the ratio between nGCaMP6m protein levels of the nmScarlet x nGCaMP6m cross and nTau^WT^ x nGCaMP6m cross / nTau^P301L,V337M^ x nGCaMP6m respectively. GCaMP6m protein quantification is shown in figure S9C and the corresponding full-length Western blots in figure S9F. Every dot represents the neuronal GCaMP6m fluorescence intensity normalized to GCaMP6m protein levels of a single animal of nmScarlet (grey) and nTau^WT^ (green) and nTau^P301L,V337M^ (orange). n = 4-5 and N = 34-37 animals. Significance was assessed by one-way ANOVA + Bonferroni post hoc test (**** = p ≤ 0.0001). E. Analysis of neurodegeneration of mechanosensory neurons as assessed by neuronal integrity of control animals and those expressing Tau^WT^ and Tau^P301L,V337M^ using *mec-4::gfp* as reporter. Images on the right show representative images and depict the individual channels and the merge. Protrusions and neurite swellings as signs of neurodegeneration were only observed for Tau^P301L,V337M^ animals (white arrows). Scale bar is 20 µm.

## Discussion

Amyloid formation is a hallmark of neurodegenerative diseases such as AD that is characterized by the deposition of Aβ plaques and Tau neurofibrillary tangles in different brain areas. We have recently established an Aβ AD *C. elegans* model that recapitulated the pathological features of AD such as aggregation, spreading and propagation of Aβ, as well as severe physiological defects such as shortened lifespan, reduced number of progenies, and impaired motility ^11^.

In this study, we have established a new *C. elegans* Tau AD model that expresses human Tau as wild-type (Tau^WT^) or mutant variant (Tau^P301L,V337M^) to complement the Aβ AD model. Previously published *C. elegans* Tau models relied mainly on the expression of untagged Tau or used a stoichiometric GFP fusion to Tau ^22,24–28^. Expression of untagged Tau maintains the characteristics of mutant variants of Tau, yet requires extraction and fractionation procedures to assess the aggregation status of Tau ^22,24–26^. Here, we used a sub-stoichiometric labelling approach that we previously used for Aβ _11_ to obtain transgenic Tau lines that express an excess of untagged Tau to preserve the properties of the protein and a sub-stoichiometrically mScarlet-tagged Tau to monitor the aggregation by FLIM in living animals. We observed an increased aggregation of Tau^P301L,V337M^ with aging and did not detect a specific onset of aggregation as previously detected for Aβ whose aggregation starts in the cholinergic IL2 neurons ^11^.

The expression and aggregation of Tau^P301L,V337M^ elicited multiple disease-associated phenotypes such as reduced lifespan, delayed development as well as impaired fecundity, motility and neuronal activity. While proteotoxicity of mutant Tau has been observed previously ^22,24–28^, this study comprehensively analyzed the aggregation of Tau^P301L,V337M^ and its detrimental effect on neuronal activity with aging. Of note, the expression of human Tau^WT^ negatively affected lifespan, fecundity and motility despite the absence of any aggregation of Tau^WT^ as reported previously ^21,27^. We cannot rule out that human Tau^WT^ binds to microtubules in C*. elegans* and thus interferes with the physiology of the animal.

Using the Aβ and Tau^P301L,V337M^ *C. elegans* AD models, we could show that neuronal activity declines already in young adult animals and thus early in the AD pathology for both Aβ and mutant Tau. The loss of neuronal activity, as assessed by GCaMP recording, manifests also in an impaired neuronal function as reflected in chemotactic defects. Although the aggregation of Aβ _11_ and mutant Tau further increases as the animals age, the neuronal activity is already at its minimum for young adult animals of these AD strains. We thus speculate that an early loss of neuronal function is a common hallmark of neurodegenerative diseases.

In contrast to studies conducted in murine APP/Aβ models, we did not detect an early hyperactivity preceding the degeneration of neurons. It is possible that other factors such as the processing of the amyloid precursor protein and genetic risk factors that are absent in *C. elegans* are required for the hyperexcitability in selected brain regions. We have, however, analyzed systemic neuronal activity in the nematode. Thus, the activity of selected neurons or neuronal circuits might differ. To address this caveat, the GCaMP sensor could be expressed in neuronal subpopulations.

Surprisingly, we observed that the expression and aggregation of Aβ in muscle tissue had a similar detrimental effect on neuronal activity as its expression in neurons. This poses the question how the proteotoxic stress in muscle tissue affects neuronal activity. Aβ can propagate from cell to cell and hence Aβ expression and aggregation in muscles can lead to spreading of Aβ into neurons ^11^. Alternatively, signaling pathways could relay the proteotoxicity of the muscle tissue to the neurons. Trans- cellular signaling of proteotoxicity and capacity of molecular chaperones has been observed before but mainly for signaling from neurons to peripheral tissue ^29,30^. Much less is known about a retrograde signaling from peripheral tissues to the nervous system. Such a signaling circuit from muscle to neurons has been observed in Drosophila. E.g., muscle FOXO/4E-BP signaling leads to a decreased feeding behavior, resulting in reduced insulin release that in turn promotes FOXO/4E-BP activity in other tissues to mitigate systemic aging ^31^. It is possible that the reduced neuronal activity in response to Aβ aggregation in muscle tissue could be communicated by a signaling pathway that may also involve neuromuscular junctions that provide a direct contact between muscle cells and neurons.

In this study, we have furthermore advanced the methodological portfolio to assess neuronal activity in living nematodes in a non-invasive manner using a microfluidic device. Neuronal GCaMP recordings require an immobilization of the animals without using anesthetics. Thus far, a myriad of microfluidic tools has been introduced that aim at supporting the investigations of *C. elegans* ^32,33^ and microfluidic devices capable of immobilizing *C. elegans* without the need for invasive, chemical paralysis have been most highly sought after. These include the on-chip use of temperature-sensitive gels _34_, quake valves and deformable membranes that collapse and compress the nematode ^35,36^, or suction channels that trap the nematodes near a side wall ^37,38^. However, unfortunately, many of these approaches rely on additional specialized equipment, thus reducing accessibility, or solely focus on the immobilization of individual specimens ^39,40^. Hence, we have decided to use passive immobilization. In contrast to previous approaches ^41,42^, our device is specifically designed for manual operation and builds on straight channels with constant widths followed by a constriction rather than tapered channels, as *C. elegans* have been observed pushing themselves off angled side walls without the constant application of a vacuum. Further, given its simple application, our device even permits the release of the nematodes post imaging, thus enabling investigations throughout their lifespan.

## Methods

### Cloning of pPD95_77::rgef-1p::GCaMP6m and mScarlet-Tau-IRES constructs

The pan-neuronal expression of the GCaMP6m was achieved by amplifying the *GCaMP6m* sequence from the pN1-GCaMP6m-XC (Addgene #111543) plasmid with following primers: 5’-CGACGACGACGACGGCTAGCATGGGTTCTCATCATCATCATCATCAT-3’ and 5’- ACGGGCGCGAGATGCGGCCGCTCACTTCGCTGTCATCATTTGTACAAACTC-3’.

The sequence of the pan-neuronal promotor *rgef-1* promoter was amplified of pPD95- DBN(wt)-YFP 17 with following primers: 5’-ACACTGCAGCATGCAAGACTAATTTTCG-3’ and 5’-ATAGGATCCGTCGTCGTCGTCGAT-3’ and cloned into the backbone of pPD95_77 (Addgene #1495) (pPD95_77::*rgef-1p)*. The GCaMP6m PCR product and the pPD95_77::*rgef-1p* were digested with NheI and NotI and Gibson Assembly (NEB) was carried out to obtain pPD95_77::*rgef-1p::GCaMP6m*.

The construct for neuronal expression of Tau^WT^ with sub-stoichiometric expression of mScarlet::Tau^WT^ and of Tau^P301L,V337M^ mScarlet::Tau^P301L,V337M^ sub-stoichiometric expression of mScarlet::Tau^P301L,V337M^ were generated by assembling 4 amplified fragments through Gibson Assembly (NEB).

The human *Tau^WT^* gene was amplified from the pRK5-EGFP-Tau (a gift from Mark Hipp, UMC Groningen, NL) plasmid with the primers 5’- GACGACCGGGATGGCTGAGCCCCGCCAGGAG-3’ with homologous overlaps to the *rgef-1* promotor sequence of the plasmid backbone at the 5’-end and 5’- AGCAACCGGTTCACAAACCCTGCTTGGCCAG-3’ with homologous overlaps to the *hsp-3^IRES^*sequence at the 3’-end. The *hsp-3^IRES^-mScarlet* sequence was amplified from the *pPD95_rgef-1p::SigPep-Aβ1−42-IRES-mScarlet-Aβ1−42::unc-54(3′UTR)*^11^ with the primers 5’-GGGTTTGTGAACCGGTTGCTCTCCCTCAC-3’ with homologous overlaps to the *Tau^WT^* sequence at the 5’-end and with 5’- GCTCAGCGGAGCTAGCCTTGTAGAGCTCGTC-3’ with homologous overlaps to the *Tau^WT^* sequence at the 3’-end. The human *Tau^WT^* gene for the expression of the Tau^WT^ sub-stoichiometrically tagged to the mScarlet fluorophore was amplified from the pRK5-EGFP-Tau plasmid with the primers 5’-CAAGGCTAGCTCCGCTGAGCCCCGCCAG-3’ with homologous overlaps to the *hsp- 3^IRES^-mScarlet* sequence at the 3’-end and with 5’- GTATCTCGAGTCACAAACCCTGCTTGGCCAGG-3’ with homologous overlaps to the plasmid backbone at the 3’-end. The plasmid backbone containing the *rgef-1* promotor sequence was amplified from the plasmid *pPD95_rgef-1p::SigPep-Aβ1−42-IRES-mScarlet-Aβ1−42::unc-54(3′UTR)*^11^ with the primers 5’- GGGTTTGTGACTCGAGATACCCAGATCATATGAAACGGC-3’ with homologous overlaps to the *Tau^WT^ sequence* at the 5’-end and with 5’- GCTCAGCCATCCCGGTCGTCGTCGTCGT-3’ with homologous overlaps to the *Tau^WT^* sequence at the 3’-end.

The human *Tau^P301L,^*^V337M^ gene was amplified from the pN1 FLTau0N4RLM YFP (a gift from Mark Hipp, UMC Groningen, NL) plasmid with the primers 5’- GACGACCGGGATGGCTGAGCCCCGCCAG-3’ with homologous overlaps to the *rgef-1* promotor sequence of the plasmid backbone at the 5’-end and 5’- AACCGGTTTACAAACCCTGCTTGGCCAGG-3’ with homologous overlaps to the *hsp-3^IRES^* sequence at the 3’-end. The *hsp-3^IRES^-mScarlet* sequence was amplified from the *pPD95_rgef-1p::SigPep-Aβ1−42-IRES-mScarlet-Aβ1−42::unc-54(3′UTR)* ^11^ with the primers 5’- GCAGGGTTTGTAAACCGGTTGCTCTCCC-3’ with homologous overlaps to the *Tau^P301L,^*^V337M^ sequence at the 5’-end and with 5’- GCTCAGCGGAGCTAGCCTTGTAGAGCTC-3’ with homologous overlaps to the *Tau^P301L,^*^V337M^ sequence at the 3’-end. The human *Tau^P301L,V337M^* gene for the expression of the Tau^P301L,V337M^ sub-stoichiometrically tagged to the mScarlet fluorophore was amplified from the pN1 FLTau0N4RLM YFP plasmid with the primers 5’-CAAGGCTAGCTCCGCTGAGCCCCGCCAGGAG-3’ with homologous overlaps to the *hsp-3^IRES^-mScarlet* sequence at the 3’-end and with 5’- TCTCGAGCTACAAACCCTGCTTGGCCAGGG-3’ with homologous overlaps to the plasmid backbone at the 3’-end. The plasmid backbone containing the *rgef-1* promotor sequence was amplified from the plasmid *pPD95_rgef-1p::SigPep-Aβ1−42-IRES-mScarlet-Aβ1−42::unc-54(3′UTR)*^11^ with the primers 5’- GCTCAGCCATCCCGGTCGTCGTCGTCGT-3’ with homologous overlaps to the *Tau^P301L,V337M^*sequence at the 5’-end and with 5’- GCTCAGCCATCCCGGTCGTCGTCGTCGT-3’ with homologous overlaps to the *Tau^P301L,V337M^* sequence at the 3’-end.

Gibson Assembly was performed following the manufacturer’s instructions to create the final plasmids that contain all 4 fragments. (*pPD95_rgef-1p::Tau^WT^-IRES- wrmScarlet-Tau^WT^::unc-54(3′UTR)* and *pPD95_rgef-1p::Tau^P301L,V337M^-IRES- wrmScarlet-Tau^P301L, V337M^::unc-54(3′UTR)*)

All cloned constructs were confirmed by sequencing (LGC Genomics, Berlin, Germany).

### C. elegans strains

All *C. elegans* strains genreated or used in this work are listed in Table 1.

**Table 1.**
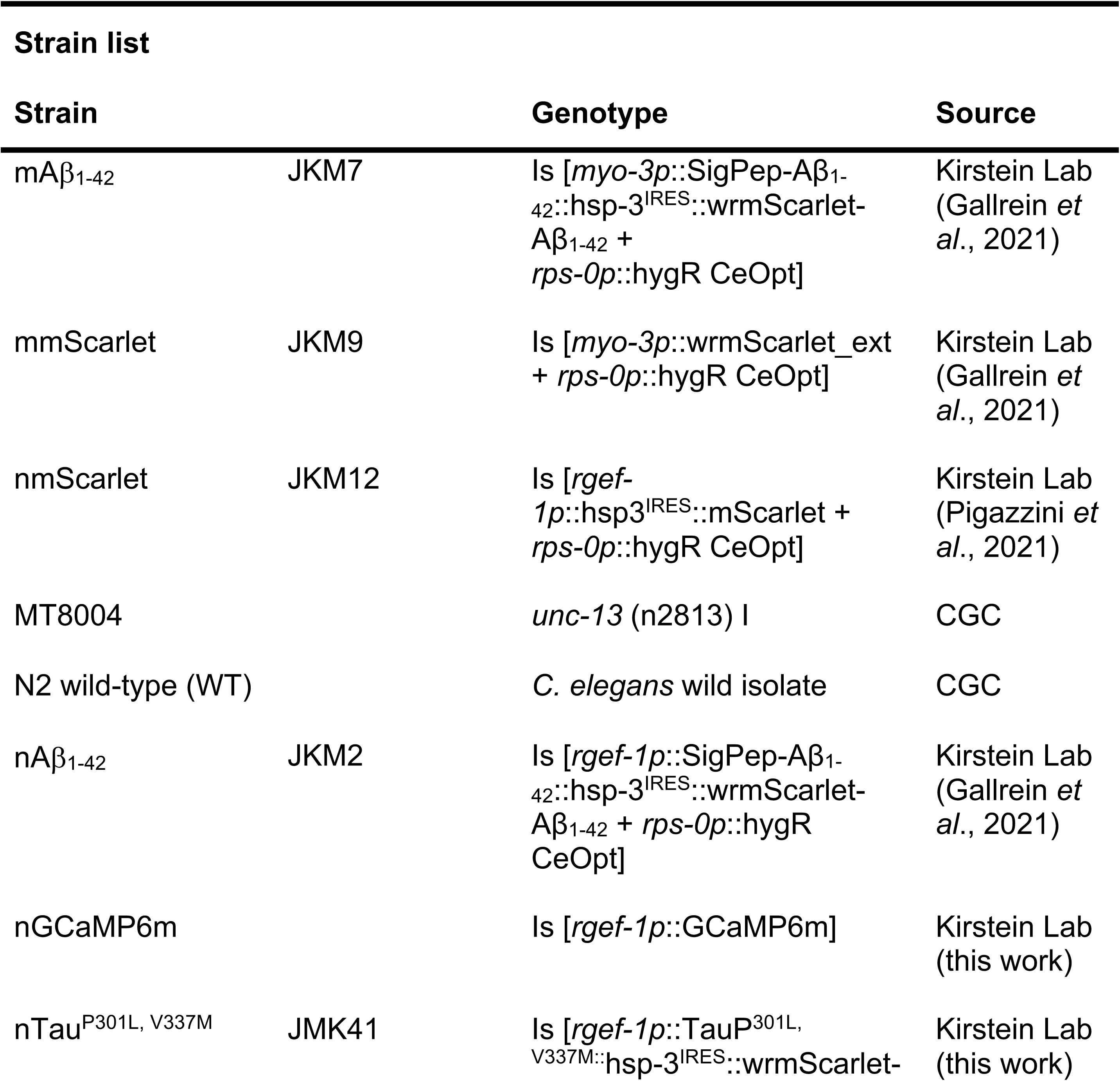
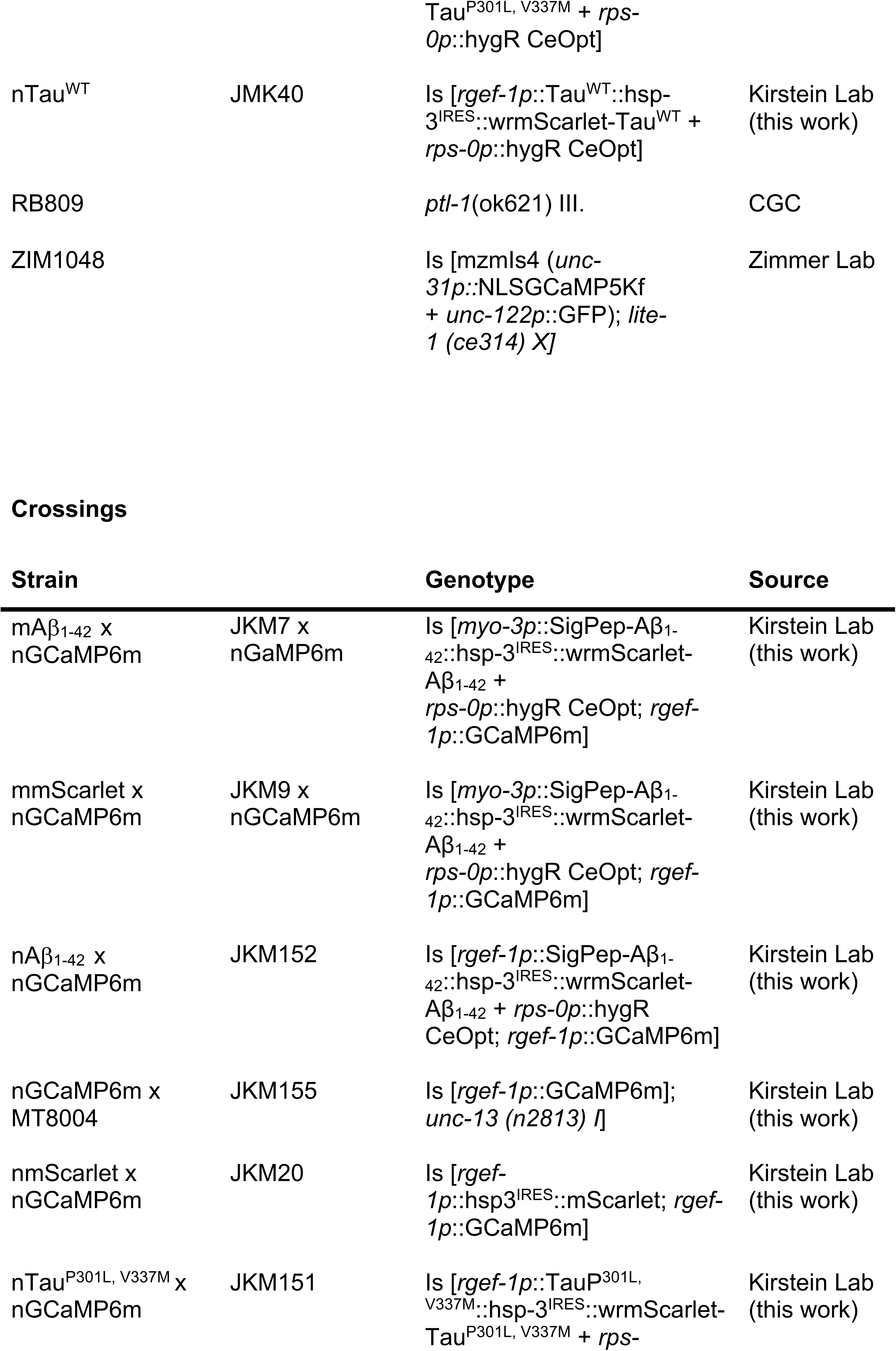
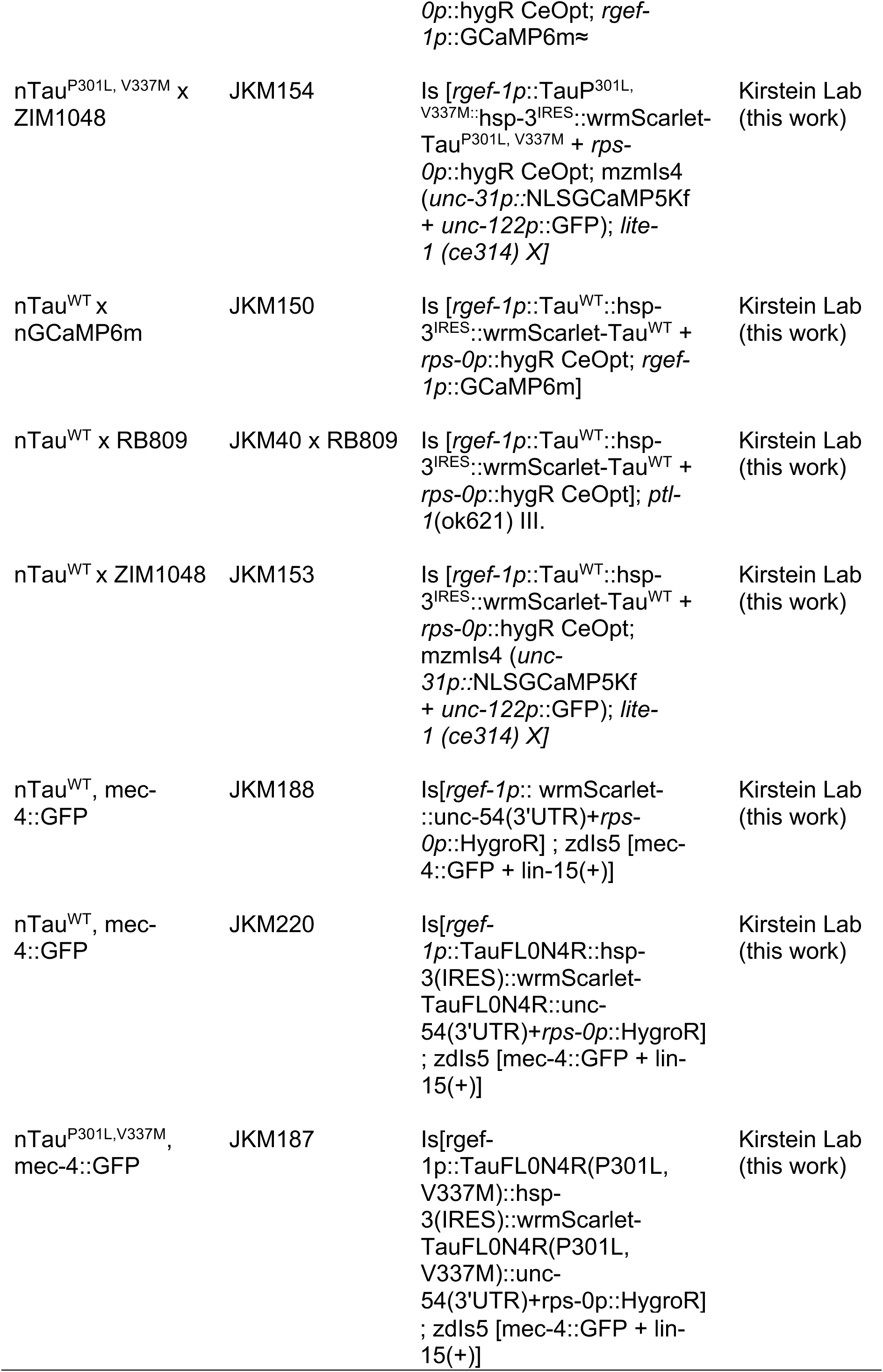
C. elegans strains used in this manuscript.

### Generation of new strains

The newly generated transgenic strains nGCaMP6m, nTau^WT^ and nTau^P301L,V337M^ have been generated by micro particle bombardment using 10 μg of the respective plasmids into N2 wild-type nematodes. Established protocols were used ^11,43,44^. Progeny was scored for red fluorescence and positive nematodes were singled out. The extra- chromosomal arrays have been integrated into the genome by UV integration. The integrated strains were backcrossed six times with N2 wild-type nematodes.

### Genetic crosses

Male nematodes used for genetic crosses were generated by exposing L4 larvae to heat treatment (31°C for 7 h). Generated males were isolated from their offspring and maintained by continuous crossing with L4 hermaphrodites. Crosses were performed by transferring L4 hermaphrodites of one genotype with an excess of males (10:1 ratio) of the other genotype on an NGM-agar plate. The desired offspring was then isolated and checked for homozygous expression of the transgenes. The offspring of the cross of the nGCaMP6m strain and the MT8004 strain was validated by genotyping with the primers 5’-TGACCACTTTGGAACCCCAT-3’ and 5’- GCATCGGAGTTTCAGTATTCTGTT-3’ using the Phire Tissue Direct PCR Master Mix kit (Thermo Fisher). The amplicons were sequenced to check for homozygosity of the single point mutation. The offspring of the strain nTauWT and RB809 was validated by genotyping using the primers 5’-CCTCCTACCACCCATCTGAA-3’ and 5’- CAACATGCTCAGGGAAGTCA-3’.

### *C. elegans* maintenance

*C. elegans* nematodes were maintained at 20°C in the dark on solid nematode growth media (NGM) seeded with live OP50 *E. coli*. All assays were performed at 20°C with age-synchronized nematodes. Young adult animals were referred to as day 4 of life. Animals were synchronized by egg-laying or by picking L4 larvae.

### RNA extraction, Reverse transcription, and real-time quantitative PCR

Nematodes were synchronised by picking 200 L4 stage nematodes of strains N2 (wild type), JKM150 (Tau^WT^), and JKM151 (Tau^P301L,V337M^). Age synchronised nematodes were grown for one, four, or seven days, rinsed off from the culture plates with M9, washed three times with M9, and resuspended in 1 ml Trizol and snap frozen in liquid nitrogen. Frozen nematode samples were thawed and lysed using three cycles of 6000 rpm shaking for 3 sec in a Precellys homogeniser. 100 µl brom-chlor-propane were added, vortexed for 15 sec and centrifuged at 12000 xg for 15 min at 4 °C. The clear phase was used for RNA extraction with RNeasy Mini Kit (QIAGEN). RNA concentration was measured with Nanodrop and RNA quality was assessed on a 1% agarose TAE gel. SuperScript® III Reverse Transcriptase (Invitrogen) kit was used for reverse transcription reactions. 1 μg of RNA was diluted in 11.5 μL total volume with RNase-free water. The mixture was heated to 70 °C for 2 min and kept on ice afterwards. 8.5 μL master mix (4 μL 5x Buffer, 2 μL DTT, 1 μL dNTP, 1 μL Oligo dT, and 0.5 μL SS III RT enzyme [Invitrogen]) was added to the RNA. The sample was incubated at 25 °C for 5 min, followed by 60 min incubation at 50 °C, and heat-inactivation at 95 °C for 15 min. 80 μL of ddH2O was added to the cDNA sample. For measuring Tau expression 10 μl PowerUp™ SYBR™ Green Master Mix was mixed with 10 mM forward and reverse primers (see below), 2 μl cDNA per sample was added and the reaction filled up with ddH2O to 20 μl. Samples were measured in triplicates. Four cohorts have been analysed. The qPCR was performed using CFX96 Touch real-time PCR detection system (BioRad). Target genes expression was normalized to expression of the reference genes (*act-1*, *lmn-1*, *eif-3.C*). The data were obtained by CFX Manager Software (BioRad) and processed with Excel 2016 (Microsoft). Primer sequences: qPCR_act-1--for 5′-CCACCATGTACCCAGGAATT-3′; qPCR_act-1--rev 5′-AGAGGGAAGCGAGGATAGAT-3′; qPCR_lmn-1--for 5′- CATCTCGTAAAGGTACTCGTAG-3′; qPCR_lmn-1--rev 5′- GTTGAGCCAAATGAATCGTC-3′; qPCR_eif-3.C--for 5′-ACACTTGACGAGCCCACCGAC-3′; qPCR_eif-3.C--rev 5′- TGCCGCTCGTTCCTTCCTGG-3′; qPCR_hTau-mScarlet-pair07--for 5′- ACCGTAAGCTCGACATCACC-3′; qPCR_hTau-mScarlet-pair07--rev 5′- GTCCCAGCGTGATCTTCCAT-3′

### Protein extraction from nematodes

Nematodes were lysed by boiling in Laemmli loading buffer. 100-125 animals of the respective age were picked into 1 ml of M9 buffer and washed three times in a low binding tube (Sarstedt). Lysis was performed in 20 μl M9 mixed with 30 μl 4X Laemmli loading buffer at 99 °C and 1000 rpm shaking for 10 min.

### Western blotting nematode lysates to quantify GCaMP6m protein levels

Nematode lysate was subjected to SDS-PAGE using an 10 % separating gel. Blotting was performed to PVDF membrane using the Trans-Blot Turbo Transfer System (Bio-Rad) and the standard program for 30 min at 25 V. Antibodies against GAPDH (Proteintech) were diluted 1:20000, against GFP(B34)/GCaMP6m (Enzo) 1:1000 and against Tau (MA5-12808, Thermo Fisher) 1:200. Proteins were detected via chemiluminescence using ECL-reagent and the ChemoStar device (Intas). Signal intensities were quantified by Fiji and normalized to GAPDH. Data are displayed as relative values to a control sample (i.e. nmScarlet, mmScarlet, nGCaMP6, nGCaMP6m DMSO treatment).

### Lifespan assay

100-180 L4 larvae were transferred in cohorts of 10-15 animals onto seeded NGM agar plates. Lifespan assay was performed as described previously ^11^. In brief, animals were transferred to fresh plates and scored daily for alive, dead or censored animals until all animals died. Animals that escaped the plates or showed a bagging phenotype were censored. Nematodes were transferred to fresh plates until day 9 to separate them from their offspring. Survival was calculated with the online tool OASIS 2 ^45^. Experiments were carried out in triplicates.

### Progeny assay

For each strain, 30-40 L4 larvae were isolated onto seeded 35 mm NGM agar plates and transferred daily onto fresh plates until they stopped laying eggs. For every nematode, the number of viable offspring was scored. Experiments were carried out in triplicates.

### Chemotaxis assay

Nematodes were synchronized and grown until the desired age and washed from the plates with M9 buffer at least five times to remove any residual bacteria. Nematodes

were kept in M9 followed by a starvation period of 90 min at 20 °C in the dark. In the meantime, unseeded NGM agar plates were marked with four quadrants and a circle in the center. Two opposite quadrants were marked as tests and 2 µl of odorant sample substances were added. The other two quadrants were marked as controls and 2 ul of control substance were added. Odorant sample substances were mixed with sodium azide (500 mM) in a 1:1 (v/v) ratio with either pure benzaldehyde (test repellent) or with diluted benzaldehyde (2 % v/v in water) (test attractant). Water was used as control substance as a mix with sodium azide. 5-10 ul of the pelleted nematodes were pipetted onto the center of the prepared plates to test around 80-100 nematodes. The plate was incubated in the dark at 20°C for 2 hours. Afterwards nematodes were counted in each quadrant and the chemotaxis index was calculated as the difference between the fraction of nematodes on the sample quadrants and on control quadrants, divided by the total number of nematodes counted. Experiments were carried out in 3-4 biological replicates and for each cohort 3 attraction and repulsion plates respectively were used.

### Developmental assay

For each strain, 60 eggs were placed onto seeded 35 mm NGM agar plates. Nematodes that did not hatch were excluded from the analysis. The start of the experiment was considered as day 1. The developmental stage of each nematode was checked daily until adulthood was reached. A nematode was considered adult when the first fertilized eggs were observed in the gonads. Experiments were performed in triplicates.

### Thrashing assay

Nematodes were analyzed in M9 media on day 4 and 10 of life. Nematodes were synchronized and grown to the desired age. 3 ml of M9 media were added to an empty 35 mm plate and 10-15 nematodes were picked into the liquid. The nematodes were allowed to swim for 3 minutes before video recording was started. Videos were recorded at 7.3x magnification at a frame rate of 30 frames per second using a stereomicroscope equipped with a camera and recording software (Leica M165 FC). Each group of nematodes was recorded three times for 20 seconds with an exposure time of 1 ms. Videos were analyzed using the WormLab software (MBF Bioscience), using the bending angle (midpoint) analysis option with an amplitude threshold of 20 degrees and a duration threshold of 10 seconds. A nematode was only included in the final analysis if the tracking lasted for at least 50% of the frames. Frequency values provided by the WormLab software were exported to Excel.

### Design and fabrication of the microfluidic mold

The design of the microfluidic channel consisted of 16 parallel microchannels which were connected to a shared inlet and outlet via regular branching (Fig. S1). Following this approach, all channels were of the same length with equal pressure loss, hence, promoting the separation of the individual nematodes after their introduction into the device. Additionally, the large number of channels permitted the distribution of the volume flow as well as the reduction of on-chip fluid velocities and, by that, allowed for a higher controllability during the loading and trapping procedure without the need for microfluidic pumps. Depending on the age of the investigated nematodes, the widths of the parallel microchannels differed between 40 μm (day 4) and 60 μm (day 7 and day 10), while the height of all channels was 65 μm. Additionally, to prevent the nematodes from getting flushed through the device during loading, each microfluidic channel was interrupted by a constriction, where its width was reduced to 12 or 20 μm, respectively. Further dimensions of the design can be found in Fig. S1A, while the corresponding lithography mask was designed in CleWin 4.0 Layout Editor (WieWeb Software) and printed on film (JD Photo Data). It is worth noting that, while tested, no pillars have been introduced near the entrance of the final design as we have not been able to confirm their previously described advantages on *C. elegans* orientation ^39^.

The microfluidic mold was fabricated using single-layer photolithography as described previously (Fig. S1C) ^46^. First, a single-side polished silicon wafer was cleaned using acetone, isopropanol, and de-ionized water, before being pre-baked at 200 °C for 10 min. Once cooled to room temperature, SU-8 50 negative photoresist (Kayaku Advanced Materials) was manually dispensed onto the wafer for spin coating. The photoresist was first distributed at 500 rpm for 15 s with an acceleration of 200 rpm/s before reduced to the final thickness using 1800 rpm for 30 s with an acceleration of 300 rpm/s. The wafer was then transferred to a hotplate for soft baking at 65 °C for 7 min before the temperature was ramped to 95 °C for a further 22 min. Finally, the wafer was cooled down to 75 °C on the hotplate before being removed and left to cool to room temperature for further processing. To transfer the design, the photoresist was exposed to 365 nm UV under contact mode for 40 s using an EVG620 NT mask aligner until a total exposure dose of 400 mJ/cm^2^ was reached. The resist was post-exposure baked at 65 °C for 1 min before being moved to another hotplate for 6 min at 95 °C and cooled down to room temperature. The design was developed in a bath with SU- 8 Developer (Kayaku Advanced Materials) for 5 min using manual agitation before development was stopped by rinsing the wafer with isopropanol followed by de-ionized water and dried using a nitrogen gun. To reduce stress inside the material, the resist was hard baked at 150 °C for 10 minutes before being cooled down to room temperature. Finally, the wafer was coated with (Tridecafluoro-1,1,2,2- tetrahydrooctyl)trichlorosilane (CAS 51851-37-7, abcr) under vacuum for 1 h to increase reusability.

### PDMS microchannel fabrication for *C. elegans* imaging

Preparation of polydimethylsiloxane (PDMS) was carried out by combining Sylgard 184 curing agent and Sylgard 184 base elastomer (Dow) in a ratio of 1:10 (w/w) followed by vigorous mixing for 5 min. The mixture was poured over the SU-8 mold (Fig. S1C) and the PDMS was degassed in a desiccator for approx. 30 min until no air bubbles were visible. The PDMS was solidified in an oven at 50 °C overnight, before being cut and carefully peeled off the wafer. The wafer sized PDMS was then separated into single devices using a razor blade. In- and outlets of 1.5 mm were applied using a biopsy punch (Ted Pella). Dust was removed from individual devices using Scotch tap before they were cleaned using soap water (Decon 90, Decon Laboratories), 70% ethanol, and ultrapure water. To remove any residual water or ethanol, PDMS devices were dried using compressed air and incubated on a hot plate at 120 °C for 5 min. Bonding of PDMS devices to glass coverslips (No. 1.5) was performed using a plasma asher (Zepto, Diener Electronic). For that, the surfaces of the glass and the PDMS were exposed to the air plasma for 30 s before being brought into contact immediately under gentle pressure. The bond was then stabilized by baking the devices at 80 °C for 5 min on a hotplate.

To load the nematodes into the PDMS channels a 1 ml low-binding pipette tip was placed at the inlet and a syringe with a flexible tubing was added at the outlet. The syringe was filled with M9 media, and the channels flushed from the outlet until the pipette at the inlet was filled half. Nematodes were picked from plates into the M9 reservoir within the pipette tip and worms were allowed to sink by gravity. By pulling slightly on the syringe, the nematodes entered the device and were separated into the microchannels where they were trapped along the sides of their bodies.

### Imaging of GCaMP6m intensity and normalization of intensities to protein levels

GCaMP6m intensities of nematodes were recorded on day 4, 7, and 10 of life, with day 1 being the day of synchronization. Nematodes were synchronized and grown until the desired age and loaded onto the fabricated PDMS microfluidic devices as described above in groups of up to six animals. For day 4-old nematodes, channels with a width of 40 um were used while older animals (day 7 and 10) were imaged in 60 um wide channels. Images of individual nematodes were taken using a custom-built widefield microscope (IX83, Olympus) with an sCMOS camera (Zyla 5.5, Andor) and a 10x objective (Plan N 10x / 0.25, Olympus). GCaMP6m was excited at 470 nm from a four-wavelength high-power light emitting diode light source (LED4D067, Thorlabs). To record GCaMP6m of individual nematodes, 2 sets of images series comprised of 10 images each were taken with an exposure time of 170 ms using the software Micro-Manager ^47^. Additionally, a brightfield image series of each nematode was taken as a reference with an exposure time of 10 ms. Fluorescence intensities of single nematodes were quantified using Fiji by selecting the most focused and stable image out of the two fluorescence image series. Normalization of the acquired GCaMP6m intensities to the GCaMP6m protein levels was performed to account for varying GCaMP6m expressions between different strains. Based on the GAPDH- normalized GCAMP6m protein levels, a normalization factor was calculated by dividing the mean expression level of control nematodes (i.e., neuronal or muscular expressing mScarlet strains) by the value for Ab/Tau-expressing nematodes for day 4, 7, and 10, respectively. Subsequently, the GCaMP6m intensity values were multiplied by the normalization factor, resulting in GCaMP6m intensities normalized to protein levels.

### Nemadipine A treatment

The L-type calcium channel inhibitor nemadipine A was dissolved in DMSO. Nemadipine A or DMSO as a solvent control was added to the liquid NGM agar at a final concentration of 2 µM before pouring. Nematodes were synchronized by egg-laying on agar plates containing nemadipine A or DMSO and grown to the desired age in the presence of nemadipine A or DMSO. nGCAMP6m intensities of nemadipine A- and DMSO-treated animals were recorded using an LSM-880 (Zeiss) confocal laser scanning microscope. Animals were immobilized on a glass slide containing a 10% agarose pad using 0.1 µm polystyrene microbeads (Polyscience 00876-15). Analysis and normalization were performed as described above.

### Confocal fluorescence imaging

Animals of a defined age were anesthetized with 250 mM NaN_3_ and mounted on a glass slide with 3% (w/v) agarose pads. Confocal images were obtained using a laser-scanning microscope LSM-880 (Zeiss). Whole nematode images were acquired using an EC Plan-Neofluar 10x/0.3 M27 objective, while head, tail, mid body and coelomocytes images were obtained using a Plan-Apochromat 40x/1,4 Oil DIC M27 objective. GFP was excited at 488 nm using an Argon laser, while mScarlet was excited at 543 nm using the HeNe laser. Transmitted-light images of the nematodes (T-PMT Transmission) were overlayed with the fluorescence image. Fluorescence intensities were quantified using Fiji by selecting nematodes individually and calculated by subtracting the product of the area selected and the average fluorescence of 3 background readings from the integrated density. Data are displayed as relative values to as control sample (i.e nTau^WT^).

### Fluorescence lifetime imaging microscopy (FLIM)

Nematodes expressing nmScarlet or mScarlet-nTau^WT^/mScarlet-nTau^P301L,V337M^ of a defined age (day 4 and 10) were anesthetized with 250 mM NaN_3_ and mounted on an 3 % agarose pad. Fluorescence lifetime imaging microscopy (FLIM) was performed using a confocal laser scanning microscope LSM-880 (Zeiss) with a pulsed laser (PicoQuant). The fluorescence lifetime was measured by time correlated single photon counting (TCSPC). mScarlet was excited with a 560 nm pulsed laser at 40 MHz with 60% laser intensity. Emission was recorded in the 575 - 625 nm range. Lifetime was monitored in the head neurons with a Plan-Apochromat 40x/1,4 Oil DIC M27 objective. Measurements were performed until the photon count reached 3000 photons in the brightest pixel using SymPhoTime 64 Software. FLIM data were analyzed using FlimFit (5.1.1) software with pixel wise fitting and assuming a mono-exponential decay. FLIM images were saved while lifetime values (τ) and histograms were exported to Excel.

## Statistical analysis

GraphPad Prism version 9 was employed for statistical analysis: Student’s t-test with Welch’s correction and One-way ANOVA with Bonferroni post hoc test were performed as parametric tests. A Two-way ANOVA was performed to test significance according to two independent variables. As non-parametric tests Mann-Whitey test was performed and Kruskal-Wallis test with Dunn post hoc test. Normality was tested by D’Agostino Pearson normality test. Significance for lifespan analysis was tested with a log-rank test using Oasis 2 online tool ^45^. P-values: ns = p > 0.05; * = p ≤ 0.05; ** = p ≤ 0.01; *** = p ≤ 0.001; **** = p ≤ 0.0001

## Acknowledgement

We thank Mark Hipp (UMCG, NL) for the Tau constructs and Olivia Masseck (Universität Köln, Germany) for the GCaMP6m construct. We appreciate the Caenorhabditis Genetics Center, which is funded by NIH Office of Research Infrastructure Programs (P40 OD010440) for the strains RB809 and MT8004. We also thank Manuel Zimmer (IMP Vienna, Austria) for the *C. elegans* strain ZIM1048. We further acknowledge funding from the Alzheimer Forschungsinitiative e.V. for the Cross Boarder grant (20052CB to JK) and a travel grant (to FH). We acknowledge funding by the University of Bremen (Fokusprojekt to JK) and the Deutsche Forschungsgemeinschaft (KI1988/7-1 to JK). G.S.K.S. acknowledges funding from the Wellcome Trust (065807/Z/01/Z, 203249/Z/16/Z), the UK Medical Research Council (MRC) (MR/K02292X/1), Alzheimer Research UK (ARUK) (ARUK-PG013-14), the Michael J Fox Foundation (16238) and Infinitus China Ltd.

---

**Figure S1.**
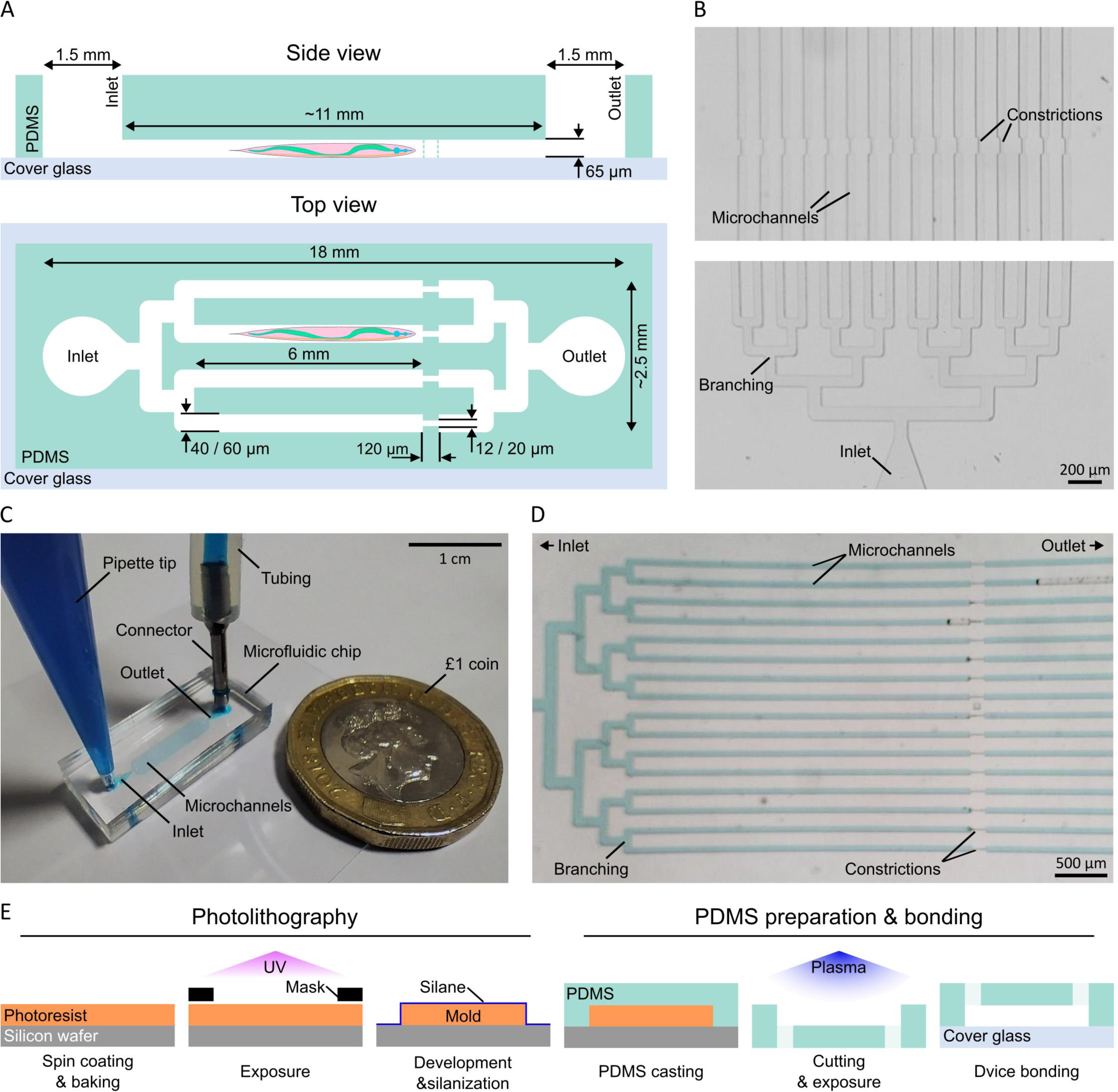
Microfluidic design and fabrication. A. Side and top view schematic of the microfluidic device. The PDMS-based device consists of 16 parallel microchannels with, depending on the age of the immobilized nematodes, widths of either 40 or 60 µm, while a uniform height of 65 µm is used throughout the device. The channels are connected to a shared inlet and outlet through regular branching. To prevent nematodes from getting flushed through the device, the channels are interrupted by constrictions. B. Brightfield microscopy images of a microfluidic device with the inlet, branching, microchannels used for trapping as well as the constrictions labelled separately. C. A photograph of the microfluidic chip next to a £1 coin for size reference. In line with the experimental procedure, a pipette tip as well as tubing have been added to the inlet and outlet, respectively. The device has been filled with blue food colouring for visualisation purposes. D. A micrograph of the same device with the 16 parallel microchannels being highlighted through food colouring. The complete area from the inlet over the branching to the constrictions is shown, which coincides with the region occupied by the *C. elegans* nematodes during the experiments. E. The fabrication procedure for the microfluidic devices can be split in two main sections, with the first one being the photolithography leading to the fabrication of the mold and the second one being the PDMS casting, preparation, and bonding. First, SU-8 photoresist is spin coated onto a wafer and exposed to UV light to create the desired pattern. Following salinization of the mold, PDMS is casted and cured, before individual chips are prepared for bonding and, finally, sealed to glass cover slips through plasma treatment.

**Figure S2.**
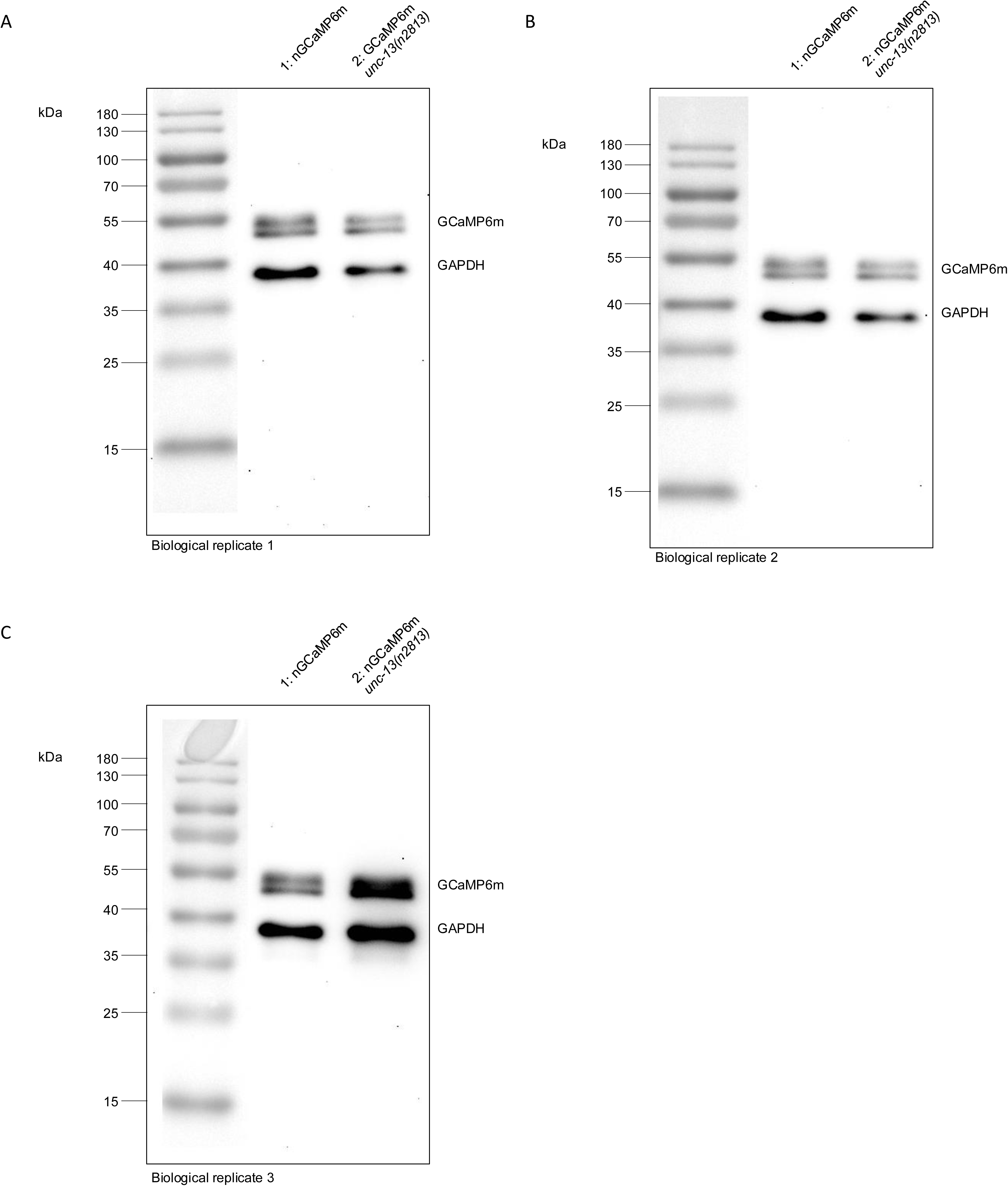
Uncropped Western blots for the quantification of GCaMP6m protein levels in nGCaMP6m and nGCaMP6m *unc-13*(n2813) animals as shown in Fig. 1E. A. Western blot of the first biological replicate of crude protein lysates of nGCaMP6m (lane 1) and nGCaMP6m *unc-13(*n2813) (lane 2) animals. Protein bands of GCaMP6m and GAPDH and molecular weights of protein ladder (kDa) are labeled. B. Western blot of the second biological replicate of crude protein lysates of nGCaMP6m (lane 1) and nGCaMP6m *unc-13*(n2813) (lane 2) animals. Protein bands of GCaMP6m and GAPDH and molecular weights of protein ladder (kDa) are labeled. C. Western blot of the third biological replicate of crude protein lysates of nGCaMP6m (lane 1) and nGCaMP6m *unc-13*(n2813) (lane 2) animals. Protein bands of GCaMP6m and GAPDH and molecular weights of protein ladder (kDa) are labeled.

**Figure S3.**
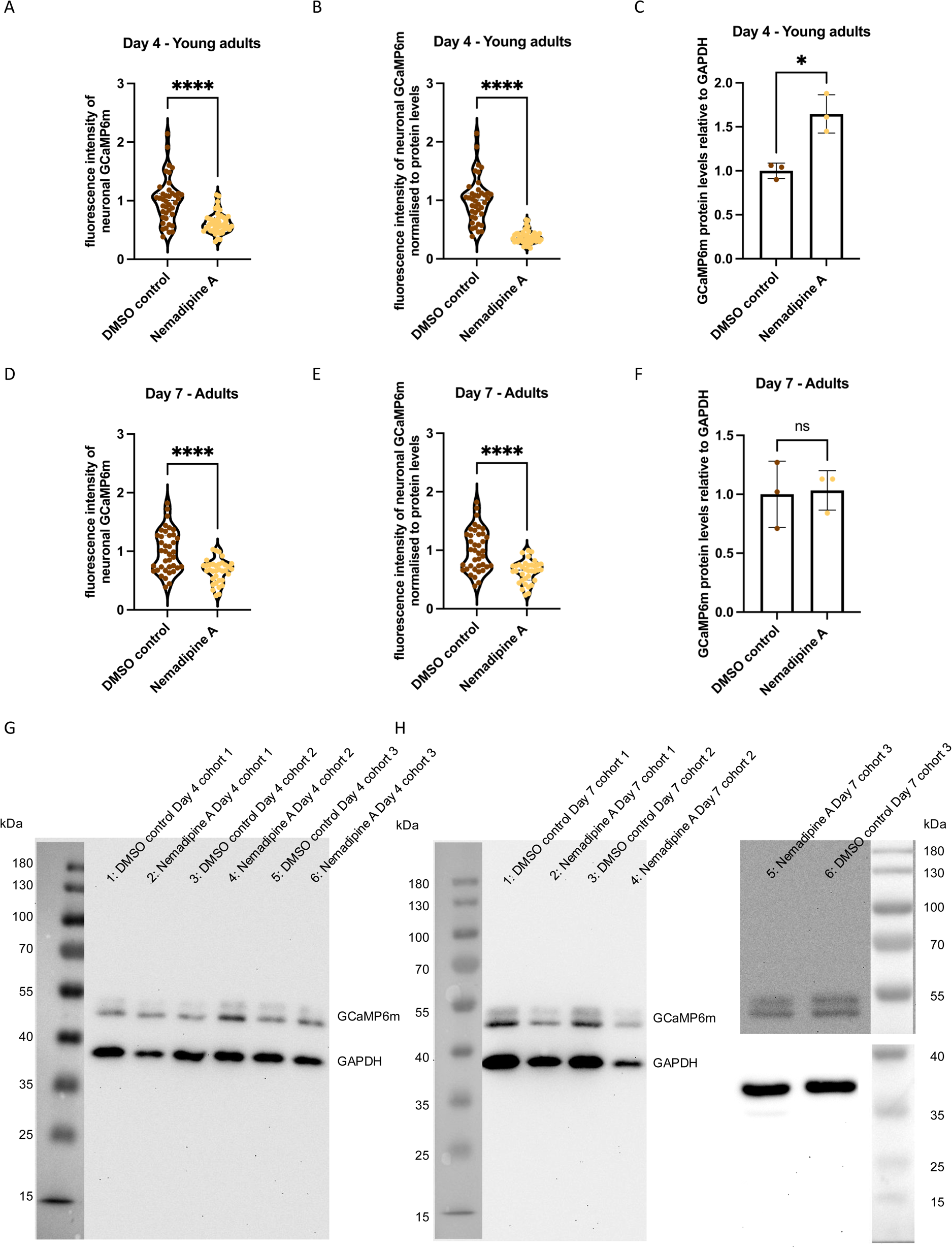
The L-type calcium channel inhibitor Nemadipine A inhibits neuronal function. A. Scatter dot plot of the average GCaMP6m fluorescence intensity of young adult animals (day 4-old) treated with Nemadipine A or with DMSO solvent control. GCaMP6m intensities were measured in alive animals immobilized with polystyrene beads with a confocal fluorescence microscope and intensities were quantified using Fiji. Every dot represents the neuronal GCaMP6m fluorescence intensity of a single animal treated with DMSO control (brown) and Nemadipine A (yellow). n = 3 and N = 40-46 animals. Significance was assessed by Mann-Whitney test (**** = p ≤ 0.0001). B. Scatter dot plot of the average GCaMP6m fluorescence intensity normalized to the GCaMP6m protein level of young adult animals (day 4-old) treated with Nemadipine A or with DMSO solvent control. GCaMP6m intensities were measured in alive animals immobilized with polystyrene beads with a confocal fluorescence microscope and intensities were quantified using Fiji. GCaMP6m intensities as depicted in S2A were multiplied with the ratio between nGCaMP6m protein levels of the DMSO treated animals and Nemadipine A treated animals. Every dot represents the neuronal GCaMP6m fluorescence intensity normalized to GCaMP6m protein levels of a single animal treated with DMSO control (brown) and Nemadipine A (yellow). n = 3 and N = 40-46 animals. Significance was assessed by Mann-Whitney test (**** = p ≤ 0.0001). C. Quantification of GCaMP6m proteins levels by Western blot from total protein lysates of young adult (day 4-old) treated with DMSO solvent control or Nemadipine A. Scatter dot plot shows quantification of GCaMP6m protein levels relative to GAPDH from three independent cohorts. Significance was assessed by unpaired Student’s t-test with Welch’s correction (* = p < 0.05). D. Scatter dot plot of the average GCaMP6m fluorescence intensity of adult animals (day 7-old) treated with Nemadipine A or with DMSO solvent control. GCaMP6m intensities were measured in alive animals immobilized with polystyrene beads with a confocal fluorescence microscope and intensities were quantified using Fiji. Every dot represents the neuronal GCaMP6m fluorescence intensity of a single animal treated with DMSO control (brown) and Nemadipine A (yellow). n = 3 and N = 35-39 animals. Significance was assessed by unpaired Student’s t-test with Welch’s correction (**** = p ≤ 0.0001). E. Scatter dot plot of the average GCaMP6m fluorescence intensity normalized to the GCaMP6m protein level of adult animals (day 7-old) treated with Nemadipine A or with DMSO solvent control. GCaMP6m intensities were measured in alive animals immobilized with polystyrene beads with a confocal fluorescence microscope and intensities were quantified using Fiji. GCaMP6m intensities as depicted in figure S1D were multiplied with the ratio between nGCaMP6m protein levels of the DMSO treated animals and Nemadipine A treated animals. Every dot represents the neuronal GCaMP6m fluorescence intensity normalized to GCaMP6m protein levels of a single animal treated with DMSO control (brown) and Nemadipine A (yellow). n = 3 and N = 35-39 animals. Significance was assessed by unpaired Student’s t-test with Welch’s correction (**** = p ≤ 0.0001). F. Quantification of GCaMP6m proteins levels by Western blot from total protein lysates of adult animals (day 7-old) treated with DMSO solvent control or Nemadipine A. Scatter dot plot shows quantification of GCaMP6m protein levels relative to GAPDH from three independent cohorts. Significance was assessed by unpaired Student’s t-test with Welch’s correction (ns = p > 0.05) G. Western blot image of crude protein lysates from three biological replicates of young adult (day 4-old) nGCaMP6m animas treated with DMSO control (lane 1, 3, 5) and Nemadipine A (lane 2, 4, 6) animals respectively. Protein bands of GCaMP6m and GAPDH and molecular weights of protein ladder (kDa) are labeled. H. Western blot image of crude protein lysates from 3 biological replicates of adult (day 7-old) nGCaMP6m animas treated with DMSO control (lane 1, 3, 6) and Nemadipine A (lane 2, 4, 5) animals respectively. Protein bands of GCaMP6m and GAPDH and molecular weights of protein ladder (kDa) are labeled. Samples of cohort 3 were analyzed on a separate gel, thus those protein bands are depicted separately from cohorts 1 and 2.

**Figure S4.**
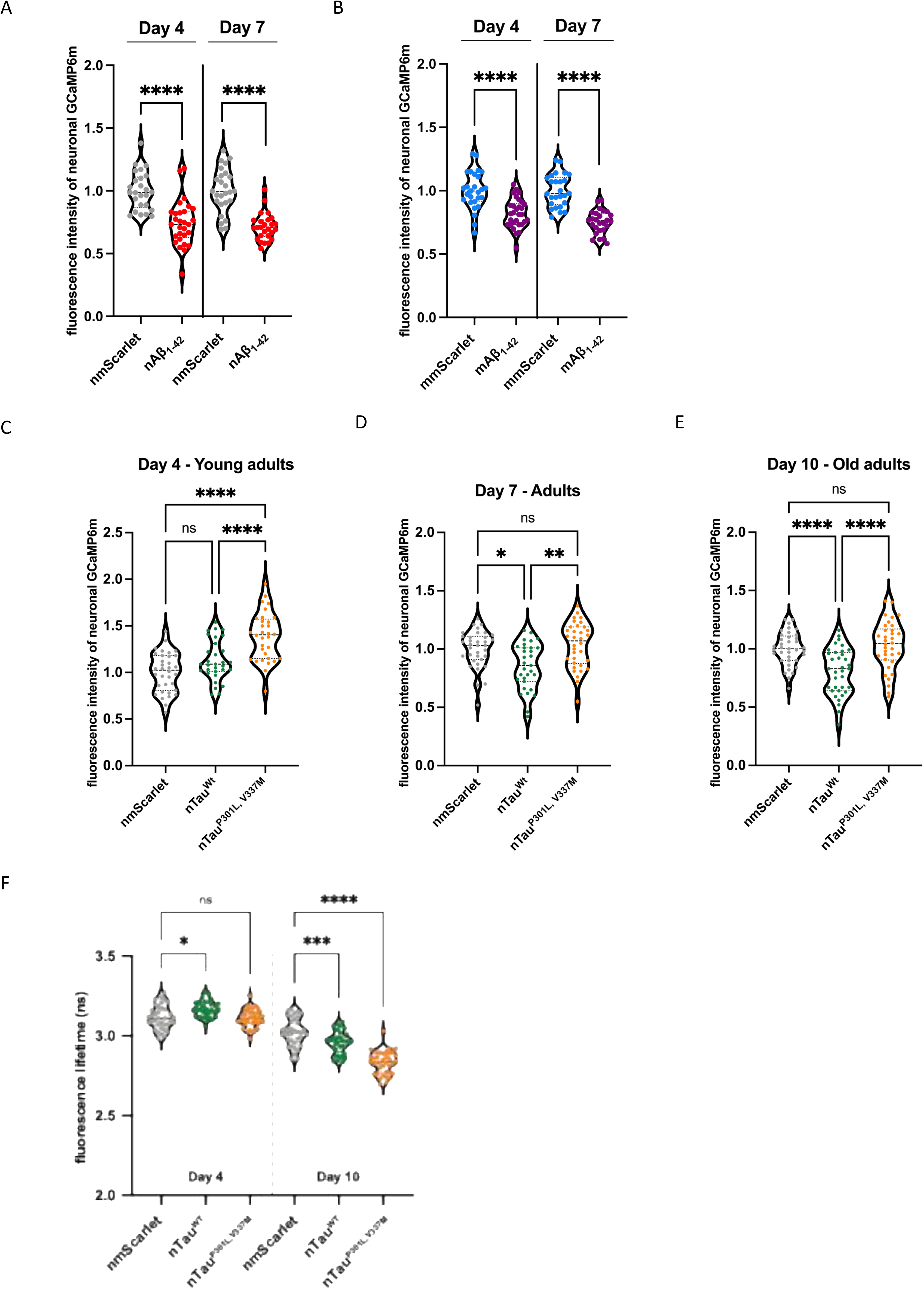
Fluorescence intensity of neuronal GCaMP6m before normalization to GCaMP6m protein levels. A. Violine dot plot of the average GCaMP6m fluorescence intensity of young adult animals (day 4-old) and adult animals (day 7-old) of nmScarlet and nAβ_1-42_ before normalization to GCaMP6m protein levels. GCaMP6m intensities were measured in alive animals using microfluidic devices with a widefield fluorescence microscope and intensities were quantified using Fiji. Every dot represents the neuronal GCaMP6m fluorescence intensity of a single animal of nmScarlet (grey) and nAβ_1-42_ (red). n = 3 and N = 25-29 animals. Significance was assessed between nmScarlet Day 4 and nAβ_1-42_ Day 4 and nmScarlet Day 7 and nAβ_1-42_ Day 7 respectively by unpaired Student’s t-test with Welch’s correction (**** = p ≤ 0.0001). B. Violine dot plot of the average GCaMP6m fluorescence intensity of young adult animals (day 4-old) and adult animals (day 7-old) of mmScarlet and mAβ_1-42_ before normalization to GCaMP6m protein levels. GCaMP6m intensities were measured in alive animals using microfluidic devices with a widefield fluorescence microscope and intensities were quantified using Fiji. Every dot represents the neuronal GCaMP6m fluorescence intensity of a single animal of mmScarlet (turquoise) and mAβ_1-42_ (purple). n = 3-4 and N = 26-30 animals. Significance was assessed between mmScarlet Day 4 and mAβ_1-42_ Day 4 and mmScarlet Day 7 and mAβ_1-42_ Day 7 respectively by unpaired Student’s t-test with Welch’s correction (**** = p ≤ 0.0001). C. Violine dot plot of the average GCaMP6m fluorescence intensity of young adult animals (day 4-old) of the control strain nmScarlet, nTau^WT^ and nTau^P301L,V337M^ before normalization to GCaMP6m protein levels. GCaMP6m intensities were measured in alive animals using microfluidic devices with a widefield fluorescence microscope and intensities were quantified using Fiji. Every dot represents the neuronal GCaMP6m fluorescence intensity of a single animal of nmScarlet (grey) and nTau^WT^ (green) and nTau^P301L,V337M^ (orange). n = 5 and N = 31-34 animals. Significance was assessed by one-way ANOVA + Bonferroni post hoc test (ns = p > 0.05; **** = p ≤ 0.0001. D. Violine dot plot of the average GCaMP6m fluorescence intensity of adult animals (day 7-old) of the control strain nmScarlet, nTau^WT^ and nTau^P301L,V337M^ before normalization to GCaMP6m protein levels. GCaMP6m intensities were measured in alive animals using microfluidic devices with a widefield fluorescence microscope and intensities were quantified using Fiji. Every dot represents the neuronal GCaMP6m fluorescence intensity normalized to GCaMP6m protein levels of a single animal of nmScarlet (grey) and nTau^WT^ (green) and nTau^P301L,V337M^ (orange). n = 4-5 and N = 31-35 animals. Significance was assessed by Kruskal-Wallis test with Dunn post hoc test (ns = p > 0.05; * = p ≤ 0.05; **= p ≤ 0.01). E. Violine dot plot of the average GCaMP6m fluorescence intensity of old adult animals (day 10-old) of the control strain nmScarlet, nTau^WT^ and nTau^P301L,V337M^ before normalization to GCaMP6m protein levels. GCaMP6m intensities were measured in alive animals using microfluidic devices with a widefield fluorescence microscope and intensities were quantified using Fiji. Every dot represents the neuronal GCaMP6m fluorescence intensity normalized to GCaMP6m protein levels of a single animal of nmScarlet (grey) and nTau^WT^ (green) and nTau^P301L,V337M^ (orange). n = 4-5 and N = 34-37 animals. Significance was assessed by one-way ANOVA + Bonferroni post hoc test (ns = p > 0.05; **** = p ≤ 0.0001) F. Violine dot plot of the average fluorescent lifetime (Δ) of young adult animals (left) and old adult animals (right) of nmScarlet, nTau^WT^ and nTau^P301L,V337M^ that also express GCaMP6m on day 4 and day 10 of life. Data displays average fluorescent lifetimes ± SD of nmScarlet (grey), nTau^WT^ (green) and nTau^P301L, V337M^ (orange). Every dot represents the average fluorescent lifetime for the head neurons of one single nematode. Three independent cohorts of in total 30 nematodes were analyzed. Significance was tested by one-way ANOVA + Bonferroni post hoc test for young adult animals and by Kruskal-Walls test with Dunn post hoc test for old animals (ns = p > 0.05; *** = p ≤ 0.001; **** = p ≤ 0.0001).

**Figure S5.**
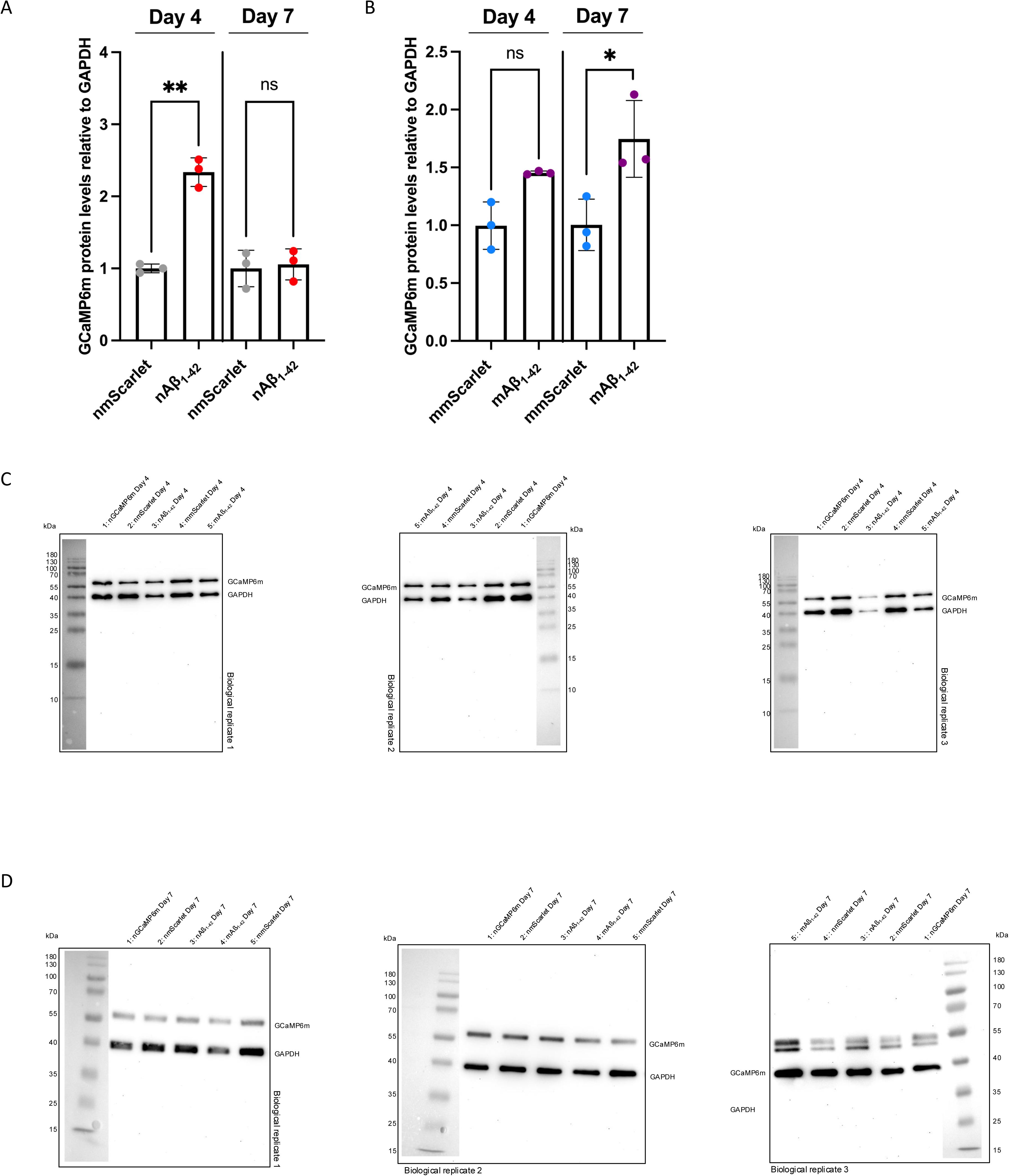
Quantification of GCaMP6m protein levels in neuronal and muscle Aβ_1-42_ and respective mScarlet control strains and corresponding uncropped Western blots. A. Quantification of GCaMP6m proteins levels by Western blots of total protein lysates of young adult (day 4-old) and adult animals (day 7-old) of nmScarlet (grey) and nAβ_1-42_ (red) animals. Scatter dot plot shows quantification of GCaMP6m protein levels relative to GAPDH from 3 independent cohorts. Significance was assessed between nmScarlet Day 4 and nAβ_1-42_ Day 4 and nmScarlet Day 7 and nAβ_1-42_ Day 7 respectively by unpaired Student’s t-test with Welch’s correction (ns = p > 0.05; **= p ≤ 0.01). B. Quantification of GCaMP6m proteins levels by Western blot of total protein lysates of young adult (day 4-old) and adult animals (day 7-old) of mmScarlet (turquoise) and mAβ_1-42_ (purple) animals. Scatter dot plot shows quantification of GCaMP6m protein levels relative to GAPDH from 3 independent cohorts. Significance was between mmScarlet Day 4 and mAβ_1-42_ Day 4 and mmScarlet Day 7 and mAβ_1-42_ Day 7 respectively by unpaired Student’s t-test with Welch’s correction (ns = p > 0.05; * = p < 0.05). C. Western blots of the first, second and third biological replicate of crude protein lysates of day 4-old nGCaMP6m (lane 1), nmScarlet (lane 2), nAβ_1-42_ (line 3), mmScarlet (line 4) and mAβ_1-42_ (line 5) animals. Protein bands of GCaMP6m and GAPDH and molecular weights of protein ladder (kDa) are labeled. D. Western blots of the first, second and third biological replicate of crude protein lysates of Day 7-old nGCaMP6m (lane 1), nmScarlet (lane 2), nAβ_1-42_ (line 3), mAβ_1- 42_ (line 4) and mmScarlet (line 5) animals. Protein bands of GCaMP6m and GAPDH and molecular weights of protein ladder (kDa) are labeled.

**Figure S6.**
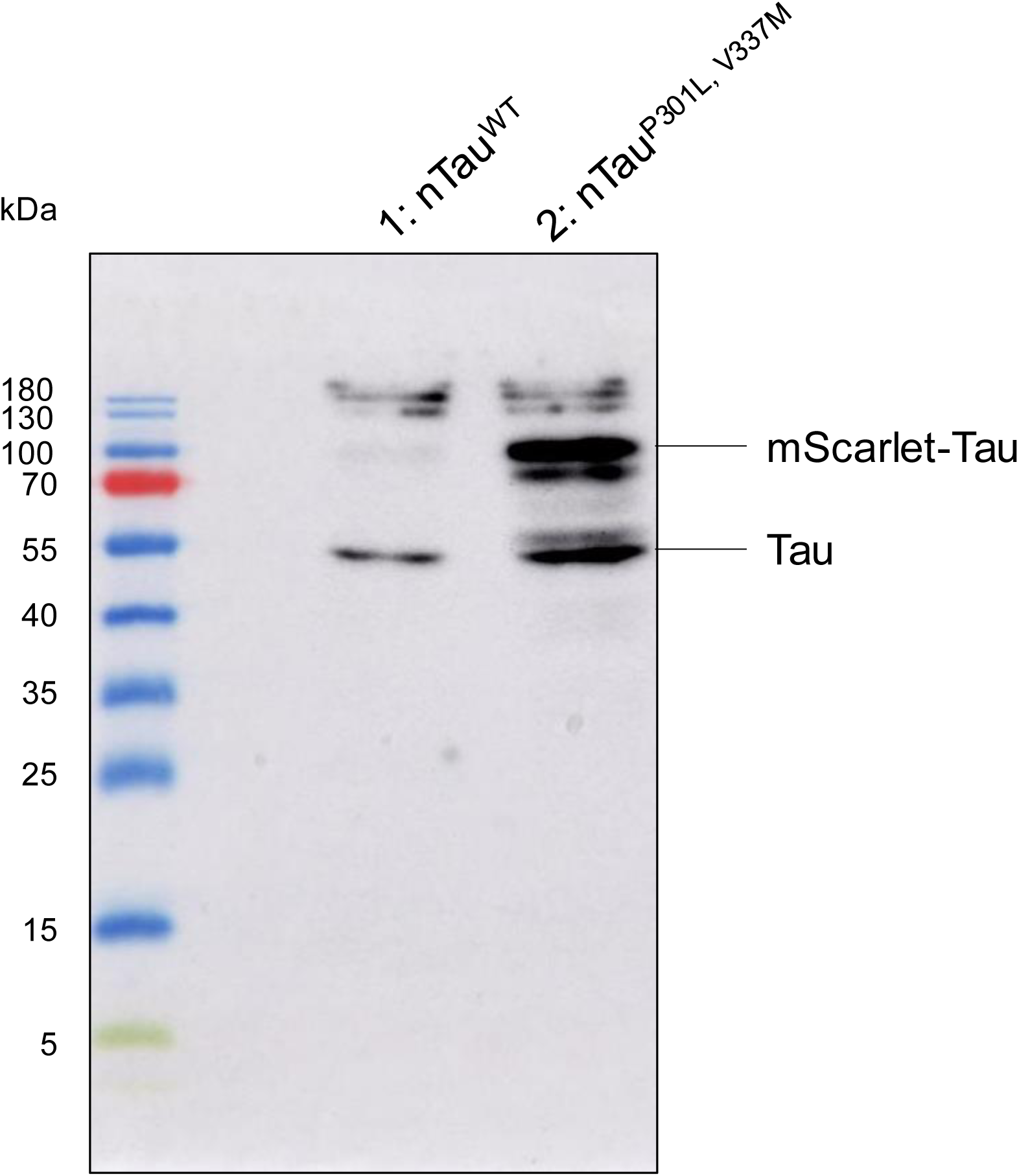
Detection of Tau and mScarlet-tagged Tau protein from lysates of nTau^WT^ and nTau^P301L,V337M^ animals. Representative uncropped Western blot of protein lysates of nTau^WT^ (lane 1) and nTau^P301L,V337M^ (lane 2). Detection was performed with Mouse-anti-Tau-5 (MA5-12808, ThermoFisher) and Goat-anti-mouse-HRP. Protein bands of Tau, mScarlet-Tau and molecular weights of protein ladder (kDa) are labeled.

**Figure S7.**
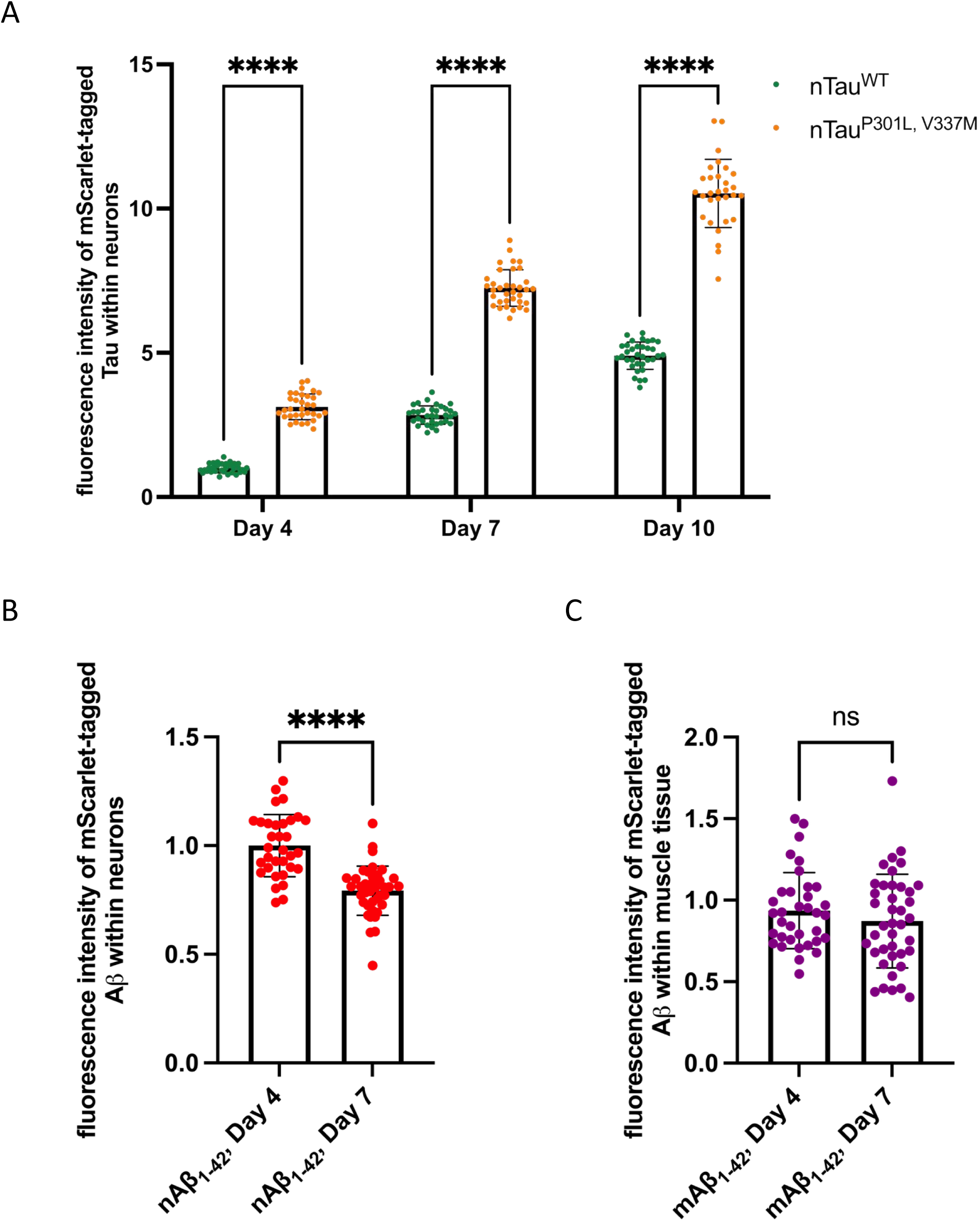
Quantification of fluorescence intensity levels of mScarlet-tagged Tau/Aβ_1-42_. A. Quantification of mScarlet-Tau^WT^ and mScarlet-Tau^P301L,V337M^ fluorescence intensity of young adult (day 4-old), adult (day 7-old) and old adult (day 10-old) nematodes of the nTau^WT^ and nTau^P301L,V337M^ strain. Confocal fluorescent images of three cohorts of total 32-35 nematodes were recorded, and fluorescence intensities were quantified by Fiji. Data are displayed as mean fluorescence intensity ± SD. Significance was tested by two-way ANOVA + Bonferroni post hoc test (**** = p < 0.0001). B. Quantification of mScarlet-Aβ_1-42_ fluorescence intensity of young adult (day 4- old) and old adult (day 10-old) nematodes of the nAβ_1-42_ strain. Confocal fluorescent images of three cohorts of total 33-38 nematodes were recorded, and fluorescence intensities were quantified by Fiji. Data are displayed as mean fluorescence intensity ± SD. Student’s t-test with Welch’s correction was performed to assess significance (**** = p < 0.0001). C. Quantification of mScarlet-Aβ_1-42_ fluorescence intensity of young adult (day 4- old) and old adult (day 10-old) nematodes of the mAβ_1-42_ strain. Confocal fluorescent images of three cohorts of total 38-40 nematodes were recorded, and fluorescence intensities were quantified by Fiji. Data are displayed as mean fluorescence intensity ± SD. Student’s t-test with Welch’s correction was performed to assess significance (ns = p > 0.05)

**Figure S8.**
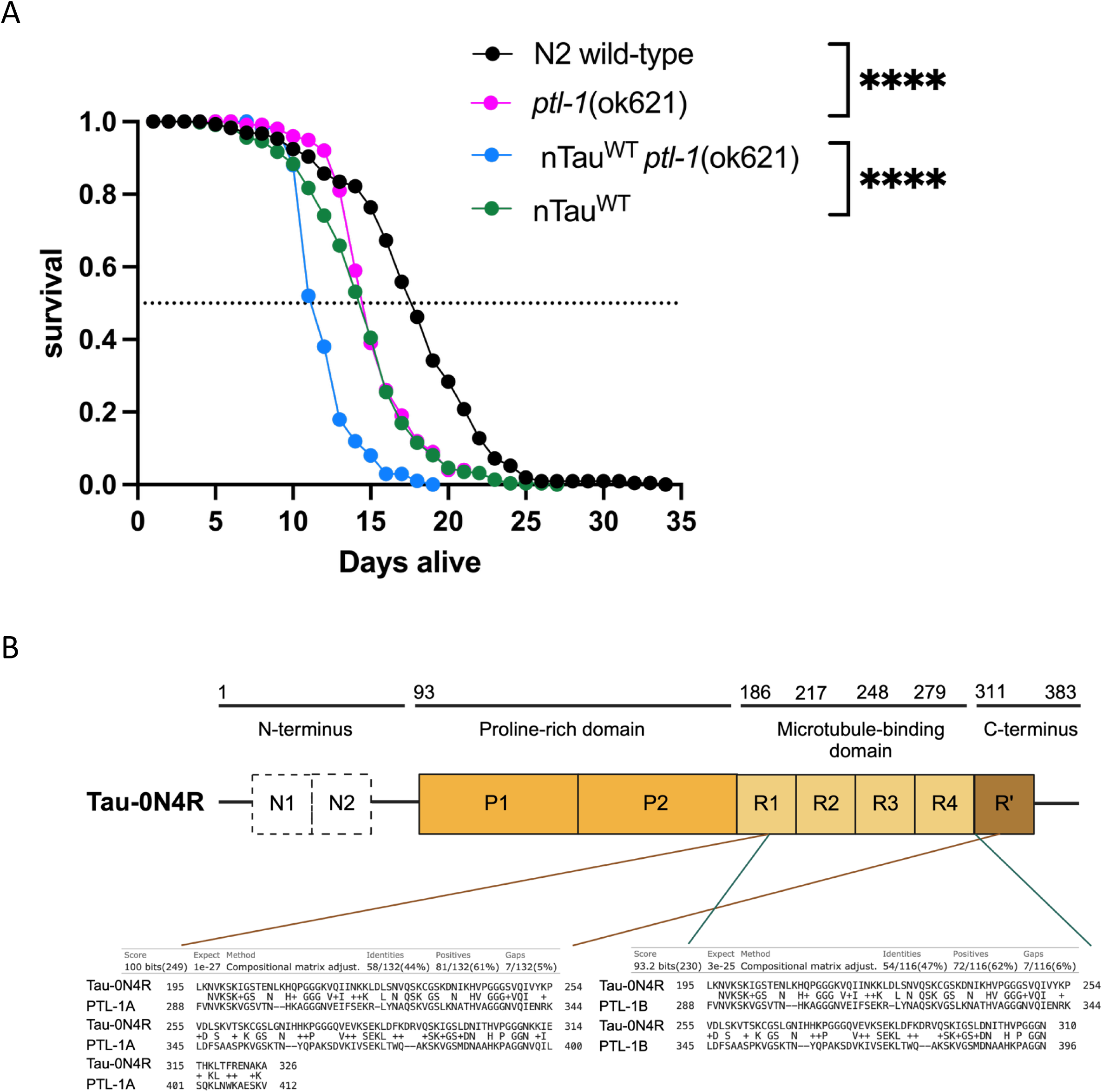
Lifespan assay of *ptl-1* knock-out mutant and nTau^WT^ cross. A Assessment of the lifespan of N2 wild-type, *ptl-1*(*ok621*), nTau^WT^ x *ptl-1*(ok621) and nTau^WT^ animals. Graph shows the cumulative survival probability (survival) versus age (days alive) of N2 wild-type (black), *ptl-1*(*ok621*) (magenta), nTau^WT^ x *ptl-1*(*ok621*) (blue) and nTau^WT^ (green). Significance was tested by log-rank test using Oasis2 online tool (**** = p ≤ 0.0001). B Protein sequences of human Tau-0N4R and both PTL-1 protein isoforms, PTL- 1A (left) and PTL-1B (right) respectively, align only within the MTBD (residues 186 to 311 of Tau-0N4R) with identities of 44 % and 47 %. NCBI reference sequences used for Protein BLAST search: NP_058518.1 (Tau 0N4R), AAB97090.1(PTL-1A) and AAC47829.1 (PTL-1B). Dashes (middle lane) denote gaps introduced into the sequences to maximise the alignment. Plus (middle lane) indicates residues that are similar but not identical.

**Figure S9.**
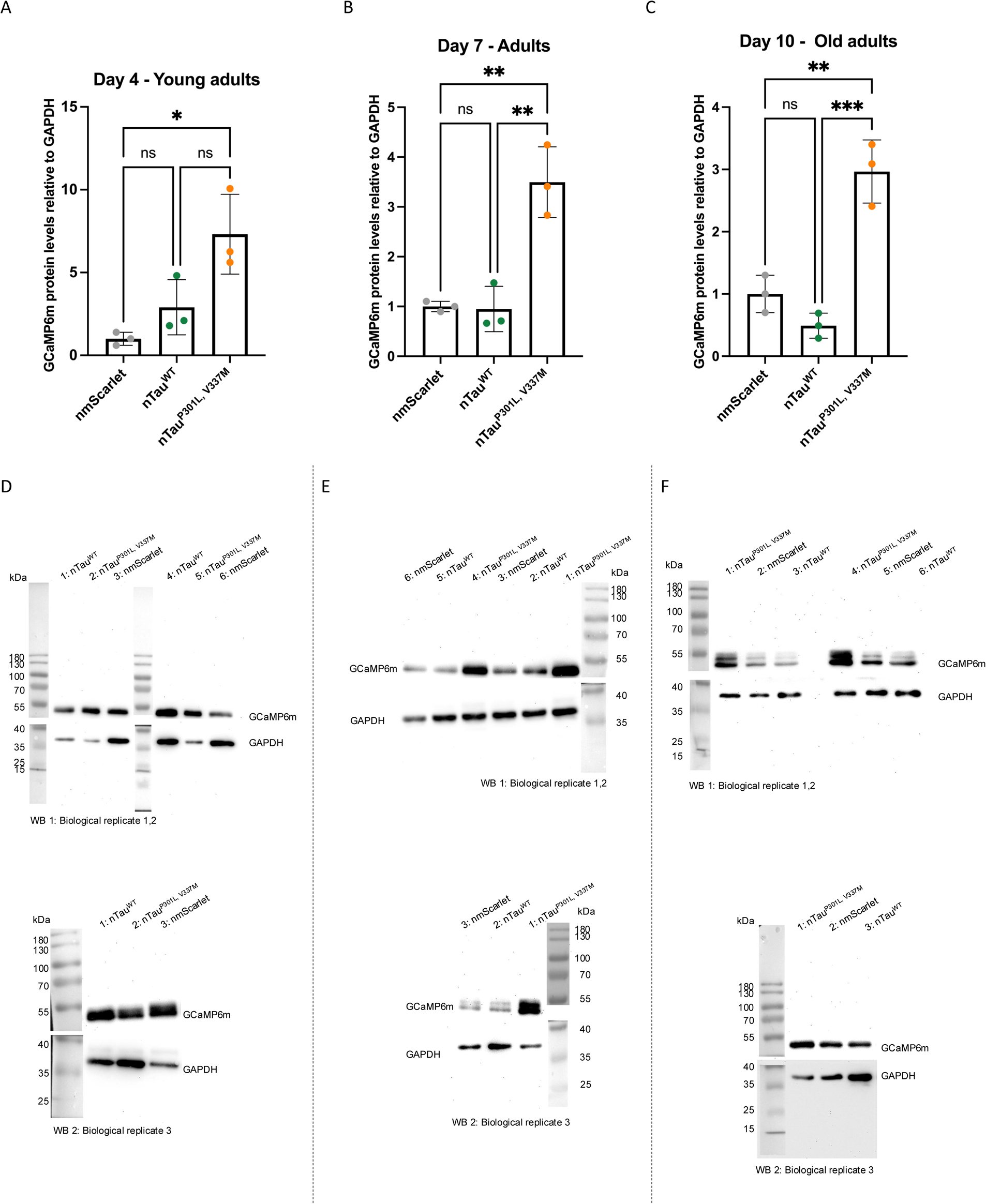
Quantification of GCaMP6m protein levels in *C. elegans* expressing Tau and nmScarlet control strain and corresponding uncropped Western blots. A. Quantification of GCaMP6m proteins levels by Western blot of total protein lysates of young adult (day 4-old) of nmScarlet (grey), nTau^WT^ (green) and nTau^P301L,V337M^ (orange) animals. Scatter dot plot shows quantification of GCaMP6m protein levels relative to GAPDH of three independent cohorts. Significance was assessed by one-way ANOVA + Bonferroni post hoc test (ns = p > 0.05; * = p < 0.05). B. Quantification of GCaMP6m proteins levels by Western blot of total protein lysates of adult (day 7-old) of nmScarlet (grey), nTau^WT^ (green) and nTau^P301L,V337M^ (orange) animals. Scatter dot plot shows quantification of GCaMP6m protein levels relative to GAPDH from three independent cohorts. Significance was assessed by one-way ANOVA + Bonferroni post hoc test (ns = p > 0.05; ** = p ≤ 0.01). C. Quantification of GCaMP6m proteins levels by Western blot of total protein lysates of old adult (day 10-old) of nmScarlet (grey), nTau^WT^ (green) and nTau^P301L,V337M^ (orange) animals. Scatter dot plot shows quantification of GCaMP6m protein levels relative to GAPDH from three independent cohorts. Significance was assessed by one-way ANOVA + Bonferroni post hoc test (ns = p > 0.05; ** = p ≤ 0.01; *** = p ≤ 0.001). D. Western blots of the first, second and third biological replicate of crude protein lysates of day 4-old nTau^WT^ (lanes 1, 4), nTau^P301L,V337M^ (lanes 2, 5), and nmScarlet (lanes 3, 6) animals. Protein bands of GCaMP6m and GAPDH and molecular weights of protein ladder (kDa) are labeled. E. Western blots of the first, second and third biological replicate of crude protein lysates of day 7-old nTau^WT^ (lane 2, 5), nTau^P301L,V337M^ (lane 1, 4), and nmScarlet (lane 3, 6) animals. Protein bands of GCaMP6m and GAPDH and molecular weights of protein ladder (kDa) are labeled. F. Western blots of the first, second and third biological replicate of crude protein lysates of day 10-old nTau^WT^ (lane 3, 6), nTau^P301L, V337M^ (lane 1, 4), and nmScarlet (lane 2, 4) animals. Protein bands of GCaMP6m and GAPDH and molecular weights of protein ladder (kDa) are labeled.

**Figure S10.**
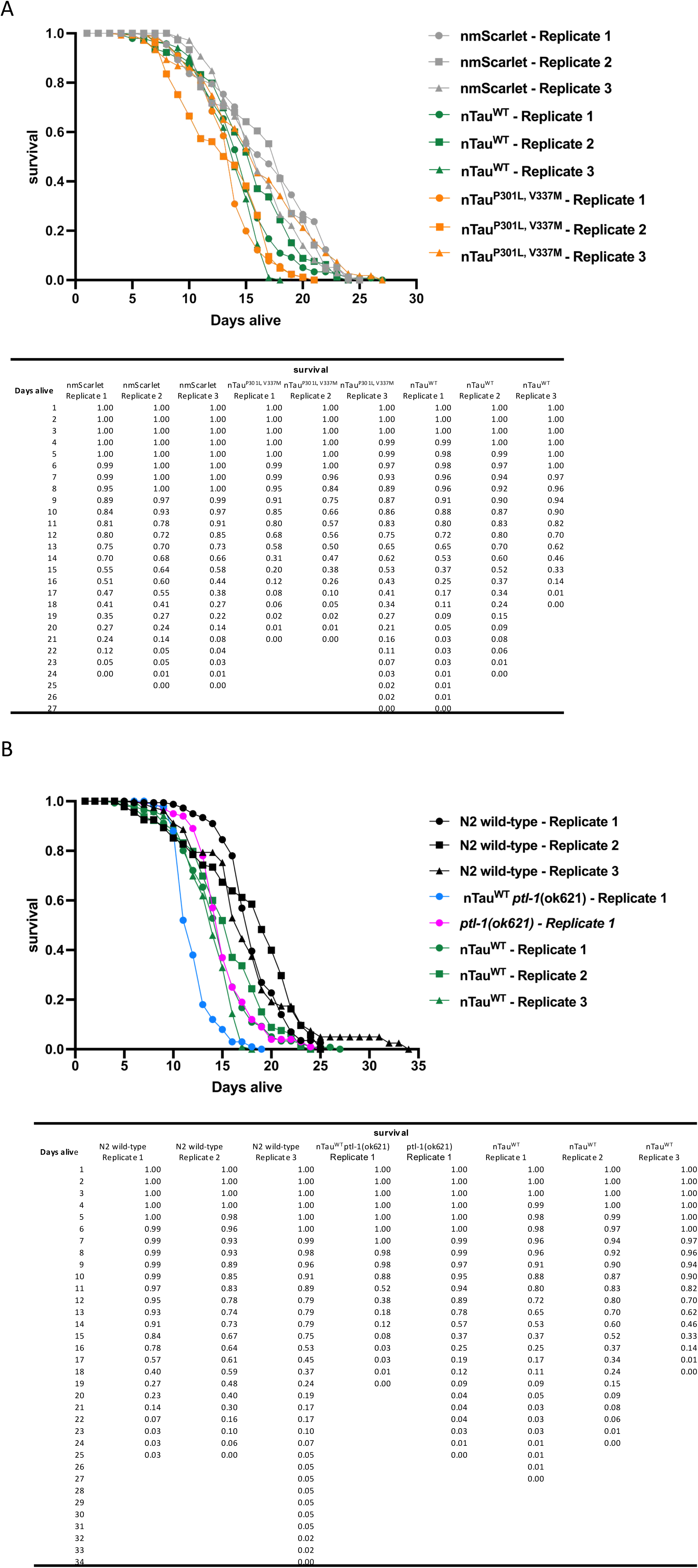
Lifespan data as graph and tabulated form. A. Lifespan data of 3 replicates each of nmScarlet (grey), nTau^WT^ (green) and nTau^P301L,V337M^ (orange) animals as graph (top) and as table (bottom). B. Lifespan data of N2 (wildtype), nTauWT, ptl-1 (ok621) and nTauWT x ptl-1 (ok621) as graph (top) and as table (bottom).

